# Modeling Synaptic Maturation from Growth Cone to Synapse in Human Organoids

**DOI:** 10.1101/2025.01.21.634055

**Authors:** Marie S. Øhlenschlæger, Lucrezia Criscuolo, Pia Jensen, Daniel J. Lloyd-Davies Sánchez, Magdalena Sutcliffe, Santosh Bhosale, Helle Bogetofte, Muhammad Tahir, Lene A. Jakobsen, Maria Pihl, Jonathan Brewer, Veit Schwämmle, Frantz R. Poulsen, Kristine Freude, Madeline A. Lancaster, Phillip J. Robinson, Martin R. Larsen

**Author notes:** Corresponding Authors: Marie S. Øhlenschlæger, Department of Biochemistry and Molecular Biology, University of Southern Denmark, Odense, Denmark, Martin R. Larsen, Department of Biochemistry and Molecular Biology, University of Southern Denmark, Odense, Denmark.

## Abstract

Human neural organoids (NOs) provide a powerful platform for investigating synaptic development and dysfunction during early neurodevelopment. However, methodologies for isolating functional synaptic structures from these models remain limited. Here, we present a differential centrifugation protocol enabling the enrichment of growth cone particles (GCPs) and immature synaptosomes from air-liquid interface cerebral organoids (ALI-COs) at distinct developmental stages (day 90 and 150). Notably, the method avoids density gradients, requires minimal starting material while maintaining reproducibility across human and murine tissues. Quantitative proteomic profiling revealed significant enrichment of growth cone markers (e.g. GAP43) and classical synaptosomal proteins (e.g. PCLO, BSN, SYN1). Transmission electron microscopy (TEM) confirmed the presence of membrane-enclosed GCPs with fibrous content and mitochondria in day 90 isolates, and immature synaptosomes containing synaptic vesicles on day 150. Functional viability of both types of synaptic structures was demonstrated through KCl-induced depolarization, which triggered phosphorylation changes in growth cone proteins (GAP43, MARCKS, MARCKSL1), cytoskeletal regulators (DCLK1, SHTN1, MARK4, MAP1B) and protein kinases (CAMK2G, PRKCE) in day 90 GCPs, as well as classical synaptic vesicle cycle proteins (SYN1, DNM1, RPH3A) at day 150. Overall, this study establishes a centrifugation-based protocol for isolating growth cones and immature synapses from human organoids, capturing key stages of synaptic development and enabling scalable, patient-compatible models to study synaptic function and dysfunction in neurodevelopmental and neurodegenerative disorders.

## Introduction

Synapses are essential for the transmission of signals between neurons. Mutations in synaptic genes have been implicated in over 130 neurological conditions, comprising both neurodevelopmental and neurodegenerative diseases (Bayés et al., 2011) and include autism spectrum disorders (Masini et al., 2020), schizophrenia (Legge et al., 2021), epilepsy (Fukata & Fukata, 2017), Alzheimer’s disease (Wang & Reddy, 2017) and Parkinson’s disease (Nguyen et al., 2019). A commonly used method to study synapses is density-based enrichment of isolated nerve terminals called synaptosomes. Synaptosomes are formed when brain tissue is homogenized in an iso-osmotic buffer, resulting in shear-induced detachment of the synaptic boutons from axons, followed by resealing of the synaptic membrane (**fig. 1A**) (Hebb & Whittaker, 1958; Whittaker, 1959). Synaptosomes contain mitochondria, synaptic vesicles, and electron dense active zone (AZ) areas with docked vesicles, which can be visualized with transmission electron microscopy (TEM). The postsynaptic density (PSD) often remains attached to the AZ areas and can be observed with TEM depending on the orientation of the section. Synaptosomes with mitochondria can be viable and metabolically active and undergo respiration in a suitable buffer (Bradford, 1970; De Belleroche & Bradford, 1972). They can regenerate their membrane potential and be depolarized by chemical stimulation with e.g., a brief pulse of elevated KCl concentration, leading to neurotransmitter release by synaptic vesicle exocytosis (Bradford, 1970; De Belleroche & Bradford, 1972; Nicholls & Sihra, 1986) followed by endocytosis to regenerate synaptic vesicles (Cousin & Robinson, 2001).

**Figure 1:**
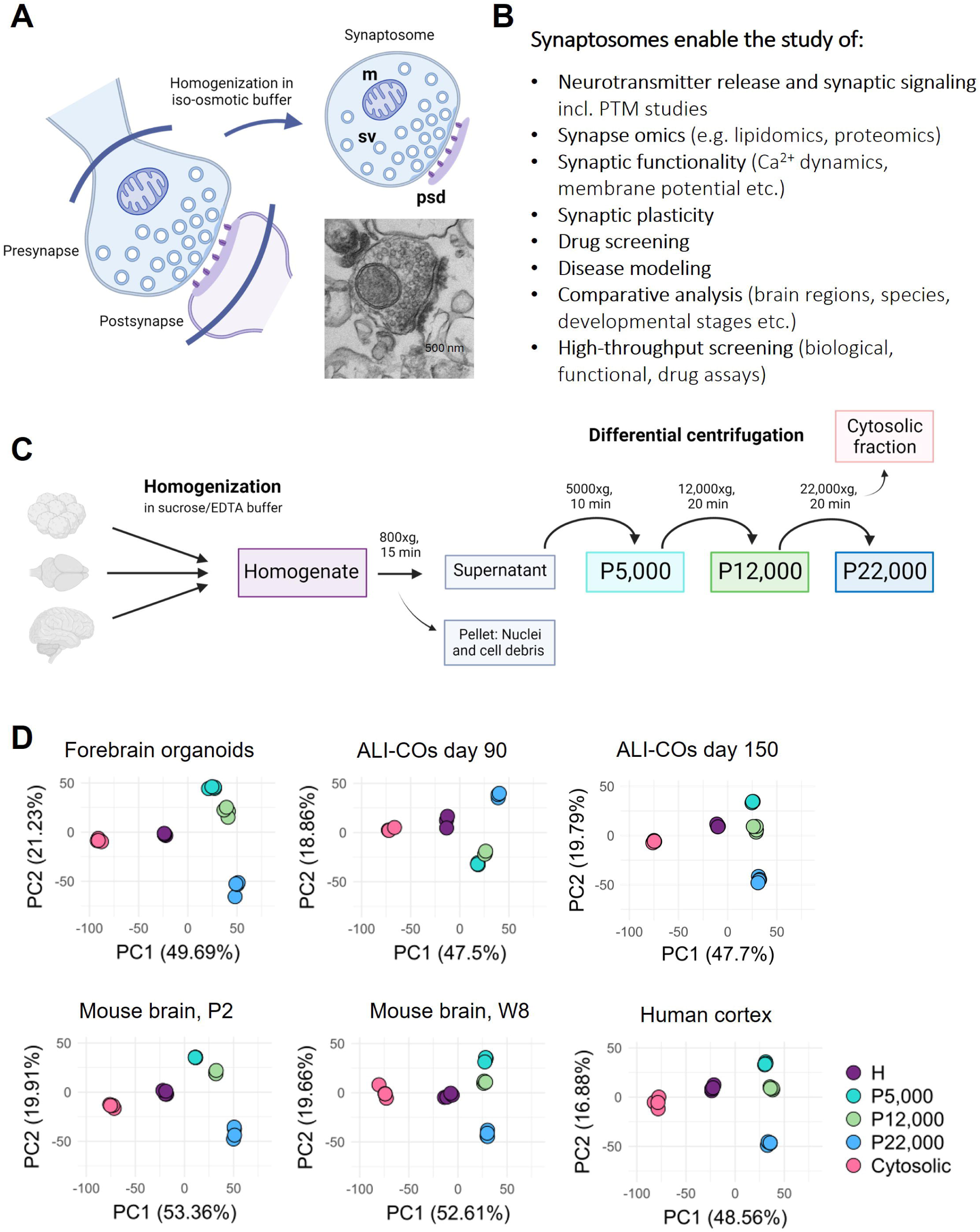
**A)** Schematic illustration of the formation of a synaptosome, including a TEM image of a mouse W8 synaptosome. m: mitochondrion, sv: synaptic vesicle, psd: postsynaptic density. **B)** List of examples of how synaptosomes can be used. PTM: Posttranslational modifications. **C)** Workflow of synaptosome enrichment using differential centrifugation. The differential centrifugation method was applied to forebrain organoids at day 100, ALI-COs at day 90, mouse brain at two developmental stages (postnatal day 2 (P2) and adult week 8 (W8)) and healthy cortical tissue from human brain. **D)** Proteomic data PCA plots showing technical replicates of the different fractions, homogenate (H), P5,000, P12,000, P22,000 and Cytosolic fraction, from differential centrifugation of human forebrain organoid tissue at day 100, ALI-COs at day 90 and 150, human cortical tissue and newborn (P2) or adult (W8) mouse brain tissue.

Traditional methods for enrichment of synaptosomes from homogenized brain tissue use different kinds of discontinuous density gradient centrifugation (Booth & Clark, 1978; Dunkley et al., 1986; Dunkley et al., 2008; Whittaker, 1959). In these protocols the brain tissue is first homogenized in an iso-osmotic sucrose/EDTA buffer followed by an initial low-force centrifugation (800-1,000xg) to pellet nuclei and non-lysed whole cells in the so-called P1 fraction. The first supernatant (S1) can then be subjected to density gradient centrifugation resulting in subcellular fractionation of the material yielding relatively purified fractions of synaptosomes. If a pure fraction is not required, the density gradient centrifugation can be replaced by a simple second centrifugation step of S1 at a higher speed, resulting in a so-called crude synaptosome fraction in the second pellet (Evans, 2015).

Synapses have a complex composition comprising several thousand different proteins involved in processes essential for synaptic function such as synaptic vesicle cycling, neurotransmitter release and synaptic plasticity (Koopmans et al., 2019). Studies of synaptosomes have played a central role in brain research with significant contributions to the present knowledge about different kinds of neurotransmitters and synaptic protein composition and function (Evans, 2015) (**fig. 1B**). Synaptosome research has benefitted from the development of mass spectrometry (MS)-based proteomic techniques that now enable reliable, large-scale identification and quantification of thousands of proteins in a single study [23, 24], together with characterization of various post-translational modifications (PTMs) of proteins, such as phosphorylation, glycosylation, cysteine modifications, acetylation and ubiquitination (Boll et al., 2020; Kang et al., 2018; Larsen et al., 2007; Leutert et al., 2021; Melo-Braga et al., 2015; Melo-Braga et al., 2014; Palmisano et al., 2012). MS-based proteomics of synaptosomes has been utilized in studies of numerous neurological disorders such as Alzheimer’s disease (Chang et al., 2013; Shen et al., 2022), schizophrenia (Paternoster et al., 2019; Zeppillo et al., 2022) and Parkinson’s disease (Betzer et al., 2015; Plum et al., 2020). The combination of techniques has been used to study stimulation-related changes in PTMs of synaptic proteins, mainly phosphorylation, providing a platform to study PTM-regulated signaling in synaptic transmission (Boll et al., 2020; Engholm-Keller et al., 2019; Kohansal-Nodehi et al., 2016; Silbern et al., 2021). Studies of depolarization-induced protein phosphorylation in synaptosomes have shown changes in phosphorylation or ubiquitination of proteins or sialylation of glycoproteins that are active in synaptic vesicle cycling, including endo- and exocytosis, actin dynamics and AZ proteins involved in neurotransmitter release (Ainatzi et al., 2025; Anggono et al., 2006; Boll et al., 2020; Engholm-Keller et al., 2019; Imoto et al., 2024; Kohansal-Nodehi et al., 2016; Silbern et al., 2021).

Synaptosome enrichment methods have traditionally been applied to rodent or human (postmortem or surgical) brain tissue. However, animal models are often not adequate when it comes to clinical translation into humans for disease mechanisms or drug development (McGonigle & Ruggeri, 2014). Synaptosome functionality from postmortem tissue is limited by the postmortem interval and often only provides information about disease endpoints, while surgical sources are rare. In extensions, synaptosomes have also been prepared from cultured neurons (Bate & Williams, 2012; Kishi et al., 1991), however with very low and crude yields. The emergence of the neural organoid (NO) field, where human pluripotent stem cells (PSCs) are differentiated in three-dimensional (3D) cultures to form brain-like tissue offers a promising new source of synapses derived from living human neuronal tissue, with potentially greater enrichment efficiency compared to 2D cultures. NOs provide a strong parallel representation of human brain *in vitro*: they recapitulate the cellular events in early human brain development (Qian et al., 2016; Renner et al., 2017) and gene expression in NOs resembles that of the human fetal brain (Amiri et al., 2018; Camp et al., 2015; Qian et al., 2016). The 3D culture of NOs promotes increased cellular interaction and cytoarchitectural organization of the neurons into cortical layers in a timed manner reminiscent of the developing brain (Renner et al., 2017). This leads to higher complexity and cellular diversity than 2D cultures. The presence of a diverse range of cell types in NOs (excitatory neurons, inhibitory neurons, intermediate progenitors, radial glia cells and astrocytes (Lancaster et al., 2013; Mariani et al., 2015; Paşca et al., 2015; Qian et al., 2016)) allows for development of functional synapses, as detected by immunohistochemical (IHC) labeling of synaptic markers and studies of electrical activity (Fair et al., 2020; Passaro & Stice, 2020; Quadrato et al., 2017; Yakoub, 2019). Spontaneous electrical activity has been recorded from NOs as early as 52-85 days of differentiation (Mariani et al., 2015; Qian et al., 2016) with later development of more coordinated network activity (3-4 months) (Fair et al., 2020) and complex dendritic morphology (day 80 of differentiation) (Giandomenico et al., 2021; Qian et al., 2016), indicative of progressive maturation of synapse physiology. Using NO tissue for synaptosome preparation could therefore provide a platform for studying synapses developed from patient-derived, induced PSCs (iPSCs), enabling functional studies of synapses in relation to diseases and drug development in a human setting. Synaptic function and synapse development could be studied in a controllable *in vitro* system that can be manipulated e.g., by CRISPR/Cas9 genetical engineering, labeling, or various kinds of stimulations.

Our aim was to establish a method for isolating viable synaptosomes from human NOs to support on-going investigations of the molecular mechanisms underlying synaptic transmission in human synapses in health and disease. The resulting method provides a straightforward, reproducible and efficient approach for enriching functional synaptic structures from NOs enabling the study of synaptic proteins during early brain development, in disease models, and in contexts such as exposure studies or drug testing.

## Methods

### Generation of dorsal forebrain organoids

Human iPSCs (IMR90-4 cells (WiCell)) were cultured on growth factor reduced Matrigel-coated (Corning) plates and daily supplemented with mTeSR1 medium (Stem Cell Technologies). The cells were used to generate dorsal forebrain organoids (FBOs) according to the STEMdiff Dorsal Forebrain Organoid Differentiation Kit (Stem Cell Technologies) with some modifications. In brief, on day 0, IMR90-4 cells were dissociated to single cells with Gentle Cell Dissociation Reagent (Stem Cell Technologies) and seeded in one well of an AggreWell 800 24-well plate at a final density of 10,000 cells per microwell (3x10^6^ cells/Aggrewell) in 2 mL FBO Formation medium supplemented with 10 μM ROCK inhibitor (Sigma-Aldrich). On day 6 embryoid bodies (EBs) were transferred to low-adherent 24-well plates with 400 μL FBO Expansion Medium until day 11 where they were moved to low-adherent 6-well plates with 2 mL medium/well and placed on an orbital shaker at 57 rpm (INFORS HT Celltron) to avoid organoid fusions. From day 43 FBOs were supplemented with 100 U/mL of Penicillin-Streptomycin (Gibco, 15140) and 1 μg/mL of Amphotericin B (Thermo Fisher Scientific). FBOs were cultured for 100 days before applying the differential centrifugation workflow. All cell and organoid cultures were maintained with 5% CO_2_ at 37°C.

### Generation of cerebral organoids and air-liquid interface cerebral organoids (ALI-CO) cultures

H9 (female) human embryonic stem cells were purchased from WiCell (WA09) and approved for use in this project by the UK Stem Cell Bank Steering Committee. Cells were maintained in StemFlex medium (Thermo Fisher Scientific) on plates coated with growth factor reduced Matrigel (Corning) and passaged twice weekly with EDTA.

Cerebral organoids were generated according to the STEMdiff Cerebral Organoid Kit (Stem Cell Technologies) seeding 2000 cells per EB. At 15 days *in vitro*, organoids were manually excised from the Matrigel droplets in which they were embedded during the kit protocol, using a needle and fine scalpel under a stereomicroscope, and returned to organoid media. From day 30, organoid media was supplemented with Matrigel (Corning) dissolved at 2% (v/v).

For culture at the air-liquid interface (ALI), cerebral organoids (COs) of day 55-60 were processed to 300 µm slices as previously described (Giandomenico et al., 2021) and collected directly onto cell culture inserts (Millipore) with serum free slice culture medium beneath (SFSCM: Neurobasal (Thermo Fisher Scientific), 1x B-27 supplement (Thermo Fisher Scientific), 1x Glutamax (Thermo Fisher Scientific), 0.5% (w/v) glucose). SFSCM was supplemented with 1x antibiotic-antimycotic (Thermo Fisher Scientific) and additional 1:1,000 (v/v) Amphotericin B (Merck, A2942-20ML). Cell culture inserts were raised vertically within their respective wells using custom-manufactured stages (MRC LMB mechanical workshop) such that more SFSCM, approximately 4.5 ml, could fit beneath each insert, permitting twice-weekly full media changes.

All cell, organoid, and ALI-CO cultures were maintained with 5% CO_2_ at 37°C.

### Mouse brains

Mouse brain tissue was obtained from 2-day-old C57BL/6J mice or 8 weeks old male C57BL/6J mice (Taconic or Janvier). Mice were housed in the Biomedical Laboratory at University of Southern Denmark (SDU) under a 12:12h light:dark cycle with food and water available ad libitum until use. C57BL/6J mice from Taconic were used for breeding mouse pups at the Biomedical Laboratory, SDU. Mice were euthanized by cervical dislocation, and their brains were immediately dissected, removing the cerebellum from adult individuals. The brains were used for either the differential centrifugation enrichment workflow (2-day-old (P2) and 8-week-old adult tissue (W8)) or a standard Percoll enrichment procedure (only adult tissue) as described below.

### Human cortical tissue

During surgery and after informed consent (approval ID S-20130048 from the Danish Research Ethics Committees), fresh human, non-tumorous cortical tissue samples were collected from a non-eloquent brain area within the exposed surgical field during surgeries for high-grade gliomas conducted at Odense University Hospital, SDU. The tissues were directly collected in ice-cold phosphate-buffered saline (PBS) and immediately processed within one hour.

### Serial differential centrifugation enrichment of synaptosomes

Mouse brains (W8 or P2), FBOs, ALI-COs or surgery-derived fresh human cortical tissue were homogenized in ice cold sucrose/EDTA buffer (0.32 M sucrose, 1 mM EDTA, 5 mM Tris base, pH 7.4, using around 700-800 µL buffer/100 mg of tissue) applying 6 up-and-down strokes with a tissue grinder (15-ml Potter-Elvehjem type Teflon-glass tissue grinder, Wheaton) at 700 rpm. A homogenate aliquot of 100-200 µL was obtained and stored at -70°C as a reference/control sample for the enrichment. The remaining homogenate was centrifuged at 800xg for 15 min at 4°C to pellet nuclei and cellular debris, and after collecting the supernatant (S1 fraction) the centrifugation was repeated after redissolving the pellet (P1 fraction) in the half amount of sucrose buffer. The protein concentration of the S1 fraction was measured using a nanophotometer (N60, Implen), and the S1 fraction was split into 3 or 4 replicates of 1.5-3 mg of protein each. Dithiothreitol (DTT) was added to the samples to a final concentration of 0.25 mM before starting the differential centrifugations (Dunkley et al., 2008). Finally, samples went through the differential centrifugations, 5,000xg for 10 min, 12,000xg for 20 min, 22,000xg for 20 min, all at 4°C, and the pellets (P5,000, P12,000, P22,000) and the supernatant from the last centrifugation (S22,000, named the “cytosolic” fraction) were collected (**fig. 1C**).

### Standard Percoll gradient enrichment of synaptosomes from mice

Mouse brains were homogenized in sucrose/EDTA buffer following a standard Percoll density gradient synaptosome enrichment procedure (Dunkley et al., 2008). Briefly, brains were homogenized with 6 up-and-down strokes at 700 rpm using a Teflon-glass tissue grinder (Wheaton) and kept on ice or at 4°C during the following procedure until stimulation. A homogenate aliquot (100-200 µL) was obtained (stored at -70°C) as a reference/control for the enrichment and the remaining homogenate was centrifuged at 800xg for 10 min, the supernatant (S1) was obtained and DTT was added to a final concentration of 0.25 mM. Carefully, 2 mL of S1 was layered on top of each Percoll density gradient (4 layers of 3%, 10%, 15% and 23% Percoll (Cytiva) in sucrose/EDTA buffer with 0.25 mM DTT) and centrifuged at 20,000 rpm for 5 min when reaching top speed in a Sorvall RC 5C plus Harvest centrifuge using a Sorvall SS34 rotor. The F3 and F4 layers were collected and washed with sucrose/EDTA buffer followed by another centrifugation at 16,000 rpm for 15 minutes. The pellets were collected into low-binding Eppendorf tubes (Sorenson BioScience Inc.), centrifuged for 10 min at 950xg and the sucrose buffer was removed. Fractions for label-free proteomic analysis were stored at -70°C.

### Depolarization of synaptosomal fractions with high KCl

Fractions from the differential centrifugation series enrichment were resuspended in 37°C HEPES-buffered Krebs-like (HBK) control buffer (4.7 mM KCl, 118 mM NaCl, 20 mM HEPES, 1.18 mM MgSO_4_, 1.2 mM CaCl_2_, 0.1 mM Na_2_HPO_4_, 10 mM glucose, 10 mM pyruvate, 25 mM NaHCO_3_, pH7.4, which had been bubbled with carbogen (95% O_2_, 5% CO_2_) for approx. 1 hour before use) and incubated at 37°C for 1 h to allow regeneration of the resting membrane potential. Subsequently, 15 sec depolarization was performed by adding an equal volume of high KCl HBK buffer (147.7 mM KCl, 20 mM HEPES, 1.18 mM MgSO_4_, 1.2 mM CaCl_2_, 0.1 mM Na_2_HPO_4_, 10 mM glucose, pH7.4) resulting in a final concentration of 76.2 mM KCl. The stimulation was stopped by snap-freezing in liquid nitrogen, and the samples were kept at -70°C until further use. Control samples were stimulated in the same way but with HBK control buffer.

### Sample lysis and digestion of proteins

The sucrose/EDTA buffer from the enrichment protocols was removed from the homogenate samples and cytosolic fractions (S22,000) using 10 kDa spin filters (Amicon Ultra-0.5 Centrifugal Filter Unit) by centrifuging at 11,000xg for 20 min at 4°C. The samples were washed x2 with 300 µL of 50 mM triethylammonium bicarbonate (TEAB), pH 8. All fraction and homogenate samples were dissolved in lysis buffer (1% sodium deoxycholate (SDC) in either 50 mM TEAB or 100 mM HEPES, pH 8) in low-binding Eppendorf tubes (Sorenson BioScience Inc.). Samples were probe sonicated 3x10 sec at 40% amplitude on ice, centrifuged for 10 min at 14,000xg and the supernatants were transferred to new tubes. Protein concentrations were measured using a Nanophotometer (Implen). An aliquot of protein (10-20 µg for label-free DIA analysis and 50-70 µg for TMT-labeling and phosphopeptide enrichment) was prepared from each sample and the proteins were reduced with 10 mM DTT at room temperature (RT) for 20 min followed by alkylation with 20 mM iodoacetamide (IAA) for 30 min in the dark at RT. After alkylation, the IAA reaction was quenched by raising the DTT concentration to 15 mM. Samples were predigested with 0.04 active units/mg Lys-C for 1h at RT followed by overnight (ON) digestion with 5% trypsin (in-house, methylated trypsin (Heissel et al., 2018)) at 37°C, pH 8. The following day, samples were incubated 1h at 37°C with 1% additional trypsin.

### Data independent acquisition (DIA) workflow using label-free quantification of proteins

After Lys-C/tryptic digestion, the SDC was precipitated using 2% formic acid (FA) followed by centrifugation and samples were lyophilized and subsequently resuspended in 0.1% FA. Peptide concentrations were measured using Pierce Quantitative Fluorometric Peptide Assay (Thermo Fisher Scientific) according to manufacturer instructions and samples were diluted accordingly in 0.1% FA to a final concentration of 0.1 µg/µL.

Differential centrifugation fractions from FBOs, mouse (P2 and W8), human cortex and Percoll density gradient F3 and F4 fractions from adult mouse brain (W8) were analyzed the following way: a total of 2 µL (200ng of peptides) was loaded onto a 15 cm x 75 µm, C18 1.6 m Aurora Elite column (ESI Source Solutions) coupled directly to a timsTOF Pro (Bruker) instrument using buffers A (0.1% FA) and B (95% acetonitrile (ACN), 0.1% FA). Peptides were eluted during a 30 min gradient (flow rate: 0.4 µL/min, increasing proportion of buffer B: from 0-2 min: 5%, from 2-18 min: 5-28%, from 18-22 min: 28-45%, from 22-23 min: 45-95%, from 23-26 min: 95%, from 26-26.5 min: 95-5%, from 26.5-30 min: 5%) on a Dionex Ultimate 3000 high performance liquid chromatography (HPLC)-system (Thermo Fisher Scientific). Samples were run scanning from 100-1700 m/z, operating in positive ion mode and using a standard DIA-PASEF (parallel accumulation serial fragmentation) method (mass range: 395.6 Da to 1020.6 Da, cycle time estimate: 1.06 sec).

The ALI-CO differential centrifugation fractions were analyzed in the same way with minor modifications in the gradient (30 min gradient at flow rate: 0.4 µL/min, increasing proportion of buffer B: from 0-2 min: 1%, from 2-2.5 min: 1-5%, from 2.5-18 min: 5-28%, from 18-22 min: 28-45%, from 22-23 min: 45-95%, from 23-26 min: 95%, from 26-26.5 min: 95-1%, from 26.5-30 min: 1%) and DIA-PASEF method (mass range: 400 Da to 1201 Da, cycle time estimate: 1.80 sec).

The mouse (P2 and W8) label-free proteomic samples were analyzed on an Orbitrap Exploris 480 mass spectrometer using a DIA method. A total of 200 ng of peptides were loaded onto a Vanquish™ Neo UHPLC system (Thermo Fisher Scientific) using a 2-column setup (Trap column of 0.5 cm, inner diameter: 300 µm and Separation column of 23 cm, inner diameter: 100 µm). Peptides were eluted with increasing amount of buffer B at a flow rate of 0.3 µL/min during a 35 min gradient: from 0-27 min: 2-29%, from 27-30 min: 29-42%, from 30-31 min: 42-70%, from 31-34 min: 70%, from 34-35: 70-100%. Peptides were analyzed using a standard DIA Scan method (precursor mass range (m/z): 400-1000, cycle time of 3 sec, orbitrap resolution: 30,000 full width half maximum (FWHM)).

### Peptide identification and label-free quantification

The raw data from DIA-MS analyses was searched with DIA-NN (v1.8.1) against computationally generated spectral libraries of human and mouse proteomes performed with the following search parameters: mass accuracy was set to 10 ppm whereas for MS1 mass accuracy to 15 ppm , peptide length range of 7-30 amino acids, precursor m/z range of 100-1700, fragment ion m/z range of 100-1700, neural network classifier in Single-pass mode, Robust LC (high precision) mode was used with retention time (RT)-dependent normalization enabled, maximum of 2 missed cleavages, N-terminal methionine (M) excision and methionine oxidation as variable modification was enabled, C carbamidomethylation was used as fixed modification.

### TMT-labeling of samples for phosphopeptide enrichment

After Lys-C/tryptic digestion, samples for phosphopeptide enrichment (high KCl stimulated fractions and controls) were labelled with isobaric Tandem Mass Tags (TMT) using TMTpro™ 16-plex or TMTpro™ 18-plex (Thermo Fisher Scientific) according to manufacturer instructions as follows: TMT-reagent (0.25 mg) for 50 µg of sample were thawed for 5 min with frequent vortexing in 100% ACN. Reagents were mixed with 50 or 70 µg of peptides per sample in lysis buffer (1:5, ACN:sample v/v). The pH was adjusted to 8 with 1 M TEAB and the labeling reaction was allowed for 1-1.5 h at RT. Labeling efficiency and TMT intensities were tested by mixing 1 µL of each sample in 2% FA followed by centrifugation at 20,000xg for 10 min to precipitate and pellet the SDC. Approximately 1 µg of the mixed sample, was run on LC-MS/MS using an orbitrap Exploris 480 mass spectrometer. The TMT labelled samples were mixed according to the obtained test. The SDC was precipitated using 2% FA and pelleted with centrifugation (20,000xg for 10 min) and the combined sample was lyophilized until a remaining volume of 100 µL, which was then used for TiO_2_ enrichment of phosphopeptides.

### TiO_2_ enrichment of phosphopeptides

The enrichment of phosphopeptides was performed on the TMT labelled peptide mixture using TiO_2_ (Larsen et al., 2007; Thingholm et al., 2006). This highly efficient method utilizes TiO_2_ enrichment in the presence of Glycolic acid, trifluoracetic acid (TFA) and ACN which dramatically prevent binding of non-phosphorylated peptides to TiO_2_ (Larsen et al., 2007). After binding of the phosphopeptides to the TiO_2_ resin (GL Sciences, Japan) in 5% TFA, 80% ACN and 1M glycolic acid for 10 min, the solution was centrifuged, and the supernatant was re-incubated with half the amount of TiO_2_ for another 10 min. The supernatant from the second incubation was collected, labelled as “non-modified” proteins and dried for further use. The TiO_2_ resin from the two incubations were washed using 200 µL loading buffer, 200 µL 80% ACN, 1% TFA and 100 µL 10% ACN, 0.5% TFA, respectively. After the final wash, the beads were lyophilized for 5 min and subsequently the phosphopeptides were eluted with 5% ammonia water, pH 11. After incubation, the supernatant was collected and lyophilized without TiO_2_ beads for subsequent high pH reversed phase fractionation.

### Desalting of “non-modified” peptides

The flow-through from the TiO_2_ enrichment was dried and subsequently resolubilized in 2 mL 0.1% TFA. The peptides were desalted on a Waters™ Oasis HLB Cartridge. The HLB cartridge was washed with 2 mL 100 % Methanol followed by 2 mL 100 % ACN using a 5 mL syringe. The HLB resin was equilibrated with 2 mL 0.1% TFA and the sample was loaded slowly using the 5 mL syringe. The HLB cartridge was washed with 2 mL 0.1% TFA and the peptides were eluted with 1 mL 50% ACN in water and subsequently the HLB cartridge was washed with 1 mL 70% ACN eluting remaining peptides. Both elutions were lyophilized and stored until further use.

### High pH reversed phase (RP) fractionation

The “non-modified” peptide and phosphopeptide samples were dissolved in high pH RP fractionation solvent A (20 mM ammonium formate, pH 9.3) and pH was adjusted with 1 M TEAB to around 9. Samples were fractionated on a Dionex Ultimate 3,000 HPLC system (Thermo Fisher Scientific) using an Acquity UPLC^®^-Class CSHTM C18 column (Waters) into 15 concatenated fractions with the following gradient of solvent B (80% ACN and 20% buffer A): sample loading with 2% solvent B, peptide elution (and fraction collection every 122 sec) from 2-50% in 59 min, 50-70% in 10 min, 70-95% in 5 min and for 10 min at 95% solvent B (during which collection of fractions was terminated). Flow rate was 5 µL/min. Fractions were lyophilized and resuspended in 0.1% FA for HPLC- Tandem MS (MS/MS) analysis.

### Data Dependent Acquisition (DDA) Nano Liquid Chromatography-Electrospray-Tandem Mass spectrometry (nLC-ESI-MS/MS)

All samples were dissolved in 0.1% FA and analyzed by nLC-ESI-MS/MS using an EASY-nLC (Thermo Fisher Scientific) with buffer A (0.1% FA) and buffer B (95% ACN, 0.1% FA) connected online to an Orbitrap Eclipse Tribrid Mass Spectrometer (Thermo Fisher Scientific, USA). The separation was performed on an in-house-made fused silica capillary two-column setup, a 3 cm pre-column (100 μm inner diameter packed with Reprosil-Pur 120 C18-AQ, 5 μm (Dr. Maisch GmbH)) and an 18 cm pulled emitter analytical column (75 μm inner diameter packed with Reprosil-Pur 120 C18-AQ, 3 μm (Dr. Maisch GmbH)). The peptides were eluted with increasing amount of the buffer B (95% ACN, 0.1% FA) from 2 to 40% in 90 and 140 min for phosphopeptide fractions and “non-modified” peptides, respectively. All spectra were generated using DDA acquisition with positive ion mode mass spectrometry. The full MS was performed in the mass range of 450–1500, in the Orbitrap with a resolution of 120,000 FWHM, a maximum injection time of 50 ms and an Automatic Gain Control (AGC) target value of 1x10^6^. Hereafter, the peptides were selected for MS/MS using higher energy collision dissociation (HCD) with normalized collision energy (NCE) setting as 35, resolution of 50,000 FWHM, AGC target value of 1x10^5^ ions and maximum injection time of 150 ms.

Additionally, the “non-modified” samples were run on the Eclipse Orbitrap in TMT Synchronous Precursor Selection Multi-Stage Mass Spectrometry (SPS-MS3) mode with real-time search (Ting et al., 2011) enabled against a Uniprot human/mouse protein database (static modifications: carbamidomethyl (C), TMTpro™ 16- or 18-plex (Kn), maximum missed cleavages: 1) with a maximum search time of 35 ms. The MS/MS isolation window was set to 2 m/z and peptides with confident identification using the built-in search engine were selected for MS3 fragmentation using HCD with NCE setting at 55, Orbitrap resolution of 30,000 FWHM, AGC target value of 500% with a scan range of 100-500 m/z.

### Peptide identification and quantification of “non-modified” peptide and phosphopeptide samples

All raw files were searched against the Uniprot database of Homo sapiens or Mus Musculus in Proteome Discoverer (PD) 2.5.0.305 (Thermo Fisher Scientific, USA) using an in-house Mascot server (v2.6) and SEQUEST HT. For HCD fragmentation, search parameters were as follows: cleavage specificity Trypsin/P; precursor mass tolerance of 10 ppm and fragment mass tolerance of 0.05 Da. The phosphopeptides search included the parameters: Maximum missed cleavages, 2; Static modifications, TMTpro (K), TMTpro (N-term) and Carbamidomethyl (C); Dynamic modifications, Phosphorylation (S, T, Y). For SPS-MS3 of “non-modified” peptides, the data were searched in PD 2.5.0.305 using SEQUEST with the same Uniprot Human/Mouse database as the real time search on the Eclipse. Search parameters were as following: cleavage specificity Trypsin/P; precursor mass tolerance of 10 ppm and fragment mass tolerance of 0.8 Da. The search included the parameters: Maximum missed cleavages, 2; Static modifications, TMTpro (K), TMTpro (N-term) and Carbamidomethyl (C). For the phosphopeptides the search was performed first in Mascot and then in SEQUEST using the same Uniprot databases and same search parameters, except for 0.05 mass tolerance in MS2 data and variable phosphorylation on S/T/Y. The percolator software in PD 2.5 was used for filtering for false discovery rate (FDR) of <1% for proteins and peptides. All datasets from PD 2.5 were exported to Excel (Microsoft) for further processing. For the Mascot searches peptides were accepted if the Mascot score was ≥15.

### Statistical analysis and data processing

Statistics of the label-free, proteomic datasets was performed with the online available PolyStest (Schwämmle et al., 2020), which does not assume normality, using the final PolyStest FDR values with Benjamini-Hochberg-based correction for multiple testing. Before statistical analysis, all label-free DIA datasets were sorted to obtain a maximum of 25% missing values for each protein across the whole dataset. Regulations in label-free datasets were considered significant when FDR ≤ 0.01 and |fold change (FC)| ≥ 1.5. For TMT-labelled phosphopeptide datasets statistics was derived from Proteome Discoverer applying the background-based T-test with Benjamini-Hochberg-based correction for multiple testing. Regulations in phosphopeptides were considered significant when adjusted P-value ≤ 0.05, fold change ≥ 1.3 up or down and grouped abundance CV < 30%. “Non-modified” TMT-labelled data was used only for identification of background protein lists for Gene Ontology (GO) term enrichment analyses.

For the proteomic DIA datasets, GO term enrichment analyses (Cellular component (CC)) were performed in Cytoscape (v3.10.0) applying the StringApp (Doncheva et al., 2019) using confidence score cutoff 0.7 and filtering for redundant terms in GO term enrichment analysis (cutoff: 0.4). Full lists of identified protein groups per dataset were used as background for each enrichment analysis, e.g., the full list of protein groups identified from the FBO fractions dataset was used as background for the GO term enrichment analysis of FBO fractions etc.

For the phosphopeptide datasets, proteins with regulated phosphopeptides from P5,000 and P12,000 fractions were analyzed together. GO enrichment analysis (CC and Molecular Function (MF)) was performed with Cytoscape (v3.10.0) and regulated phosphopeptides of two selected protein-protein interaction (PPI) clusters (confidence score cutoff 0.4) were visualized using the OmicsVisualizer app (Legeay et al., 2020). All identified master proteins in “non-modified” samples (based on ≥ 2 unique peptides) and phosphopeptide datasets combined were used as background for the enrichment analysis.

Dot plots, principal component analysis (PCA) plots and bar-graphs were made in R (v4.3.0) (R Core Team, 2013) using the package ggplot2 (Wilkinson, 2011) and ChatGPT (vAugust 3, 2023) was used, for coding assistance. Heatmaps were made using the pHeatmap package (Kolde, 2019) in R.

The statistical test used for changes in synaptic and growth cone marker proteins was a Wilcoxon Rank Sum test, suited for data where normal distribution and equal variance cannot be assumed. Tests were conducted as one-sided, testing for up-regulation in P5,000, P12,000 and P22,000 fractions and down-regulation in Cytosolic fractions.

### Immunofluorescent labeling

Organoids were fixed with 4% paraformaldehyde (Thermo Fisher Scientific) in PBS (Gibco) for 1 h at RT and soaked in 30% sucrose in PBS ON or until further processed. Organoids were embedded in OCT mounting media (VWR), snap frozen in < -50°C ethanol and sectioned in 30 µm sections on a Leica CM1860 cryostat at -17 to -20°C. Sections were placed on microscope slides (Thermo Fisher Scientific) and stored at -20°C. Slices were hydrated with PBS and washed 2x5 min with wash buffer (0.1% TritonX100 (Plusone) in PBS). Permeabilization and blocking buffer (5% donkey serum (BioWest)) was added followed by 30 min incubation at RT. Primary antibodies diluted in wash buffer incl. 1% donkey serum (BioWest) were added (see dilutions in **table 1**) and slices were incubated ON at 4°C to allow binding. Slices were washed in wash buffer for 3x10 min followed by incubation for 1 h in the dark with secondary antibodies diluted in wash buffer and 1% donkey serum (see dilutions in **table 2**). Secondary antibodies were removed, and slices were washed with PBS for 3x10 min followed by mounting with DAPI-containing ProLong Diamond Antifade mountant (Invitrogen). The slices were stored in the dark at 4°C until image acquisition. The images were obtained with confocal microscopy on a Nikon A1R confocal unit coupled to a Ti-2 LFOV microscope. Images were analyzed with ImageJ (v1.53t).

**Table 1:** Primary antibodies for labeling of synapses in FBO and ALI-CO slices.

| Antigen | Species | Company | Cat. No. | Dilution |
| --- | --- | --- | --- | --- |
| MAP2 | mouse | Sigma | M1406 | 1:2000 |
| PSD95 | goat | Abcam | ab12093 | 1:100 |
| SYP | mouse | Merck Millipore | S5768 | 1:200 |
| SYN1 | rabbit | Abcam | ab254349 | 1:500 |
| VGLUT1 | rabbit | Synaptic Systems | 135303 | 1:1000 |

**Table 2:**
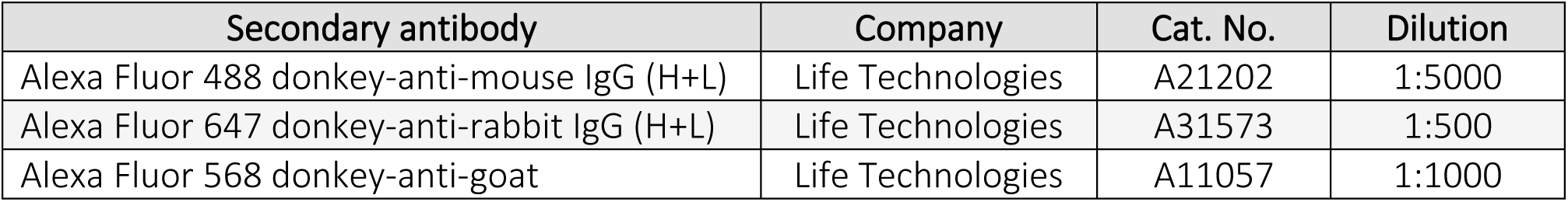
Secondary antibodies for labeling of synapses in FBO and ALI-CO slices.

| Secondary antibody | Company | Cat. No. | Dilution |
| --- | --- | --- | --- |
| Alexa Fluor 488 donkey-anti-mouse IgG (H+L) | Life Technologies | A21202 | 1:5000 |
| Alexa Fluor 647 donkey-anti-rabbit IgG (H+L) | Life Technologies | A31573 | 1:500 |
| Alexa Fluor 568 donkey-anti-goat | Life Technologies | A11057 | 1:1000 |

### Transmission electron microscopic (TEM) analysis

Fractions were fixed in 3% glutaraldehyde in 0.1 M sodium phosphate (SP) buffer (78 mM Na_2_HPO_4_, 22 mM NaH_2_PO_4_, pH 7.3) for 0.5-1 h on ice. The fixing solution was replaced with 0.1 M SP buffer and samples were stored at 4°C until further processing. The fractions were embedded in 4% Batco agar at 45°C (BD diagnostics) and washed x2 in 0.1 M SP buffer. The samples were postfixed in 1% OsO4 (in 0.1 M SP buffer) for 1 h at RT and covered in folio followed by washing x2 in Milli-Q water. The samples were subsequently dehydrated in a series of increasing concentrations of ethanol (EtOH): 50% EtOH (10 min), 70% EtOH (10 min), 96% EtOH (10 min) and 99% EtOH (3×20 min). The EtOH series was followed by an intermediate solution of propylene oxide (Merck) (2×10 min). Samples were then gradually infiltrated in Epon (812 Resin, TAAB T031) and incubated in a rotor ON in pure Epon. Next, the samples were embedded in Epon and polymerization was allowed for 48 h at 60°C. The specimens were cut into semi-thin sections of 2 μm with a glass knife (LKB Bromma 7800, Leica Microsystems) on an ultramicrotome (Reichert Ultracut S, Leica, Microsystems) followed by staining with 1% toluidine blue (VWR) in 0.1% Borex (VWR), to identify a section of interest. Ultra-thin sections of 50–60 nm were cut with a diamond knife (MD1035) on an ultramicrotome (Reichert Ultracut S, Leica, Microsystems) and collected on 150 mesh copper grids (Gilder) and coated with a parlodion-amyl acetate film (EMS). Finally, the sections were contrasted using 2% uranyl acetate (Polyscience, 21447) and 1% lead citrate (Reynold, 1963). The micrographs were examined with a Talos L120C transmission electron microscope operating at 80 kV and images were acquired using a Ceta camera and Velux software (Olympus).

### Western blotting

Samples were lysed in 1% SDC and 50 mM TEAB, pH 8, and sonicated 3x10 sec on ice at 40% amplitude. Following sonication, samples were centrifuged at 14,000xg for 10 min at RT and supernatant transferred to new Eppendorf tubes (Sorenson BioScience Inc.). Protein concentration was determined by Nanophotometer (Implen). For each sample 3.2 µg protein was separated using Bolt 4-12% Bis-Tris Plus Gels (Thermo Fisher Scientific) with samples loaded in NuPage LDS Sample buffer (Thermo Fisher Scientific) with NuPage Sample Reducing agent (Thermo Fisher Scientific) and PageRuler Plus pre-stained protein standard (Thermo Fisher Scientific) for size reference. Proteins were transferred to activated PVDF membranes (Millipore) using the Trans-Blot^®^ Turbo™ Transfer System (BioRad) and membranes were blocked with 5% skimmed milk in TBS-Tween20 (TBST) buffer for 1 h and incubated overnight at 4°C in 5% milk/TBST with the following primary antibodies: Anti-rabbit Translocase of outer mitochondrial membrane 20 (TOMM20) (Abcam) 1:1,000 or mouse anti-

Synaptophysin (SYP, Millipore) 1:200. Anti-mouse β-actin HRP-linked (Abcam) 1:50,000 was used as loading control. Following 3 x wash in TBST, the membranes were incubated for 2 h at RT in TBST with the following secondary antibodies: anti-rabbit IgG, HRP-linked (Cell Signalling) 1:10,000 or anti-mouse IgG, HRP-linked (Abcam) 1:10,000. Following 3 x wash in TBST, the membranes were visualized with Immobilon ECL Ultra Western HRP Substrate (Millipore) using an Amersham 680 Imager (GE Healthcare).

## Results

### The differential centrifugation protocol is highly reproducible across brain tissue types

After several attempts to enrich synaptosomes from NOs using a standard Percoll density gradient method yielding no clear F3/F4 (synaptosome enriched) layers, we decided to develop a different approach. Differential centrifugation strategies have previously been used in combination with TMT-based proteomics to characterize organelle-specific proteins in various cell types. A study from 2019 by Geladaki et al. observed mitochondria pelleting at 3,000xg, Peroxisomes at 5,000xg, lysosomes at 12,000xg and Golgi and ER from 15,000-30,000xg (Geladaki et al., 2019). Based on an estimated density of synaptosomes, we expected these structures to be pelleted around 5,000-12,000xg. The workflow we developed for differential centrifugation enrichment of synaptic structures from human NOs is shown in **fig. 1C**. This protocol was applied to multiple neural tissue types including two types of NOs (FBOs at day 100 and ALI-COs at day 90 and 150, cell culture images are shown in fig. S1A) together with newborn mouse brain (postnatal day 2, P2), adult (week 8, W8) mouse brain and adult human cortical tissue (surgery derived, healthy tissue). Prior to the enrichment from NOs, the presence of synapses in the NO tissues was confirmed with IHC labeling of pre- and postsynaptic markers (Synapsin-1 (SYN1), Vesicular glutamate transporter 1 (VGLUT1), synaptophysin (SYP), Postsynaptic density protein 95 (PSD95, also called DLG4), and Microtubule-associated protein 2 (MAP2)), showing co-localization in both organoid types (fig. S1B-D). A standard procedure for enrichment of synaptosomes, based on discontinuous Percoll density gradient centrifugation (Dunkley et al., 2008), was performed in parallel on adult mouse brain (W8) for comparison. The Percoll-based synaptosome enrichment generated the synaptosome-containing fractions F3 and F4 and the pellet from the first 800xg centrifugation (P1) was used as a negative reference for synapse-specific proteins for the Percoll enrichment procedure. Four technical replicates of the new differential centrifugation samples were examined with LC-MS/MS-based DIA proteomics and principal component analysis (PCA) showed similar fraction patterns between each of the tissue types, indicating high reproducibility between brain species and organoid types (**fig. 1D**).

### Evaluation of protein content from the differential centrifugation fractions

To assess enrichment across fractions, we identified fraction-specific proteins (defined as fold change (FC) ≥ 1.5 relative to the corresponding total homogenate sample, FDR < 0.01) and performed Gene Ontology Cellular Component (GO CC) enrichment using the full dataset as background. **Figure 2** summarizes enriched GO CC terms for each fraction derived from FBOs, ALI-COs (day 90 and 150), newborn mouse (P2) and adult human cortex; top20 results from each tissue are in **Supplementary fig. S2A-X**.

**Figure 2:**
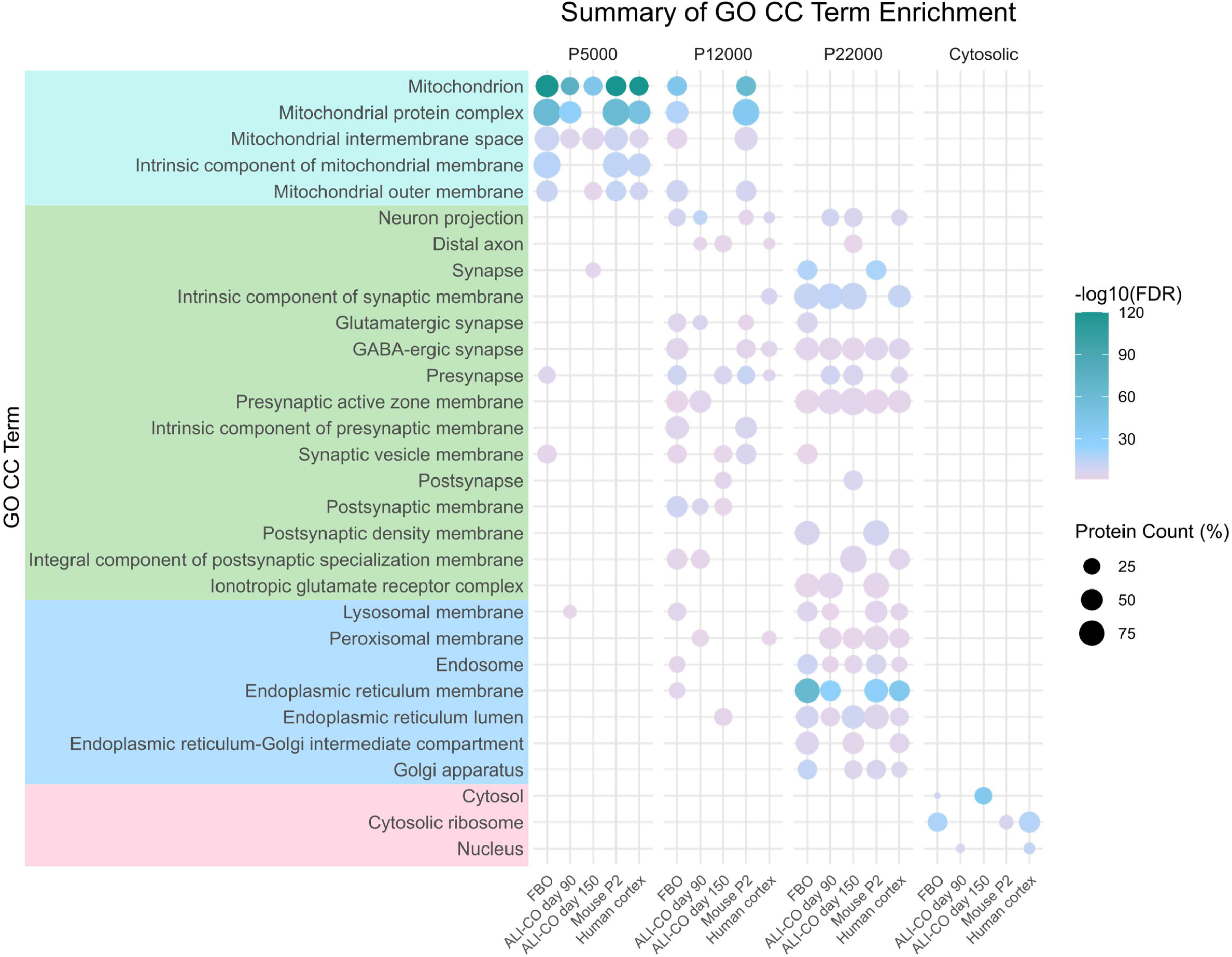
Summary of Gene Ontology Cellular Component (GO CC) enrichment in the different fractions (Homogenate (H), P5,000, P12,000, P22,000 and Cytosolic) from the differential centrifugation protocol. The enrichment was based on fraction-specific proteins (FC ≥ 1.5 compared to homogenate, FDR ≤ 0.01) for each of the five tissue types: forebrain organoids at day 100 (FBO), ALI-COs at day 90 and 150, mouse brain at postnatal day 2 (P2) and healthy cortical tissue from human brain (Human cortex). Mint: Mitochondrial terms; green: neuronal and synaptic terms; blue: endomembrane system terms; pink: cytosolic or nucleus terms. The terms were selected manually based on designated key-words (e.g. “synaptic”, “mitochondrial”, “lysosomal”, “golgi”, “cytosolic” etc.) together with the average FDR across tissue types, total number of proteins and number of times the term appeared across the datasets. All enriched term FDRs are below 0.05.

For all tissues, P5,000 fractions were enriched in mitochondrial proteins. In addition, FBOs and day 150 ALI-CO also showed enrichment of synaptic terms, suggesting pre-synaptic enrichment. In contrast, day 90 ALI-COs and human cortex P5,000 fractions primarily showed mitochondrial terms. All P12,000 fractions showed enrichment in several synaptic terms, together with mitochondrial, lysosomal, and a few ER-related terms. Synaptic terms in the FBO and in both day 90 and day 150 ALI-COs included both pre- and postsynaptic terms (**fig. 2** and **Supplementary fig. S2F and J**). Human cortex P12,000 fraction revealed synaptic and ion channel-related enrichment (**fig. 2** and **Supplementary fig. S2V**). P22,000 fractions also showed substantial enrichment in synaptic terms; however, this was accompanied by significant enrichment in endomembrane-related terms (e.g., endosome, ER and Golgi). Mitochondrial terms were absent. Cytosolic fractions were enriched in cytosolic and nuclear proteins, consistent with partial nuclear disruption during homogenization.

Overall, the GO enrichment patterns were highly consistent across the tissue types investigated, suggesting similar organelle distributions between organoids and human/mouse brain tissue. This was supported by Western blot analysis of the mitochondrial marker TOMM20, which confirmed declining mitochondrial distribution across fractions from FBO and human cortex (**Supplementary fig. S3A**).

### Enrichment of synaptic proteins by differential centrifugation

To delve into the GO enrichment patterns we compared synaptic protein enrichment in organoid fractions to adult mouse brain tissue (W8) processed with traditional Percoll density gradients. We focused on a panel of 30 synaptic proteins (**Supplementary table S1**), quantified across all datasets (25 detected in ALI-CO fractions). Their abundance in the pellet fractions (P5,000, P12,000, and P22,000) was compared to the corresponding total homogenate sample levels, using the corresponding Percoll gradient method fractions (F3 and F4) as positive controls (**fig. 3A**).

**Figure 3:**
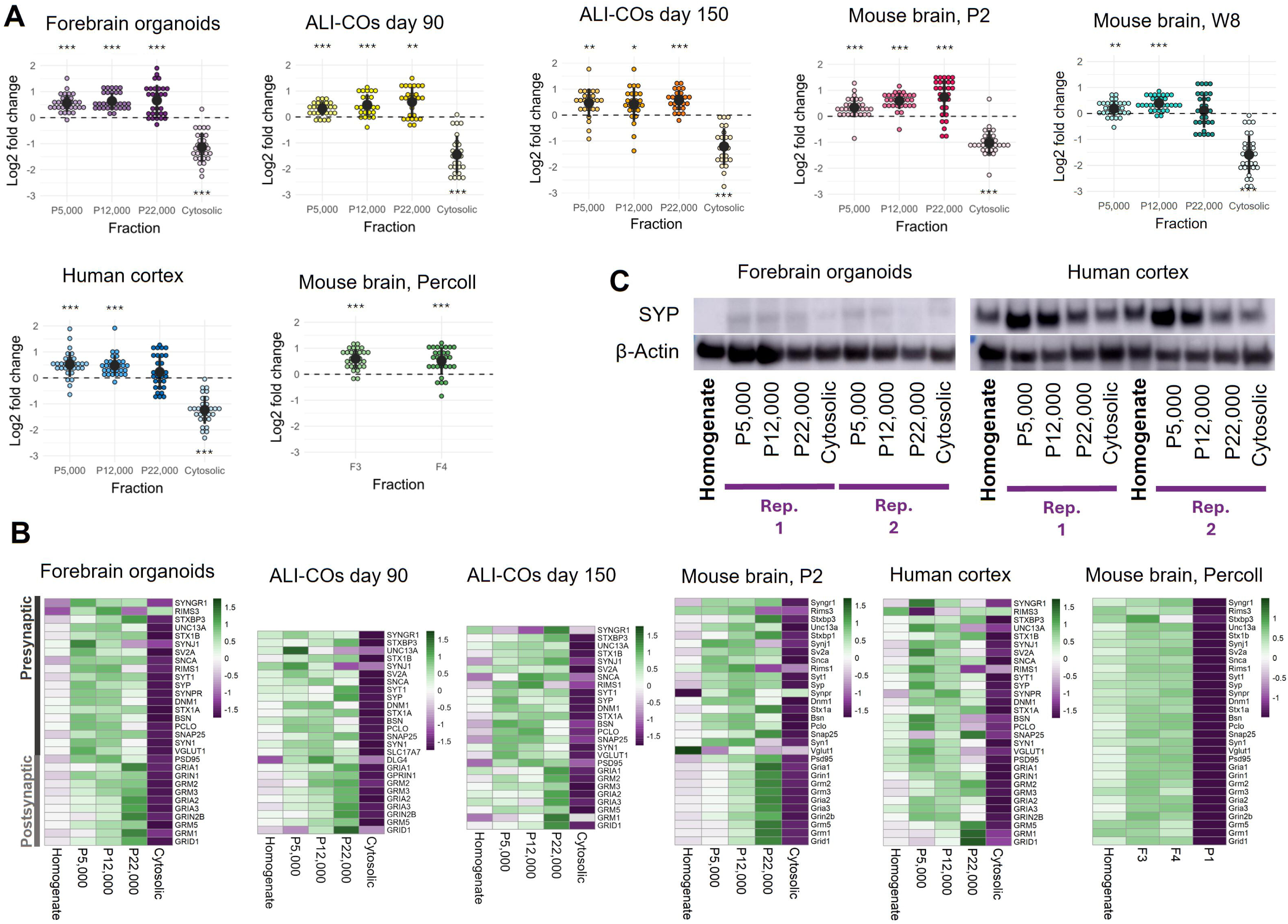
**A)** Changes in the levels of 30 selected synaptic markers in the differential centrifugation fractions compared to the corresponding total homogenate samples from each tissue type. An enrichment of synaptosomes from adult mouse brain (W8) using the conventional Percoll^®^ density gradient centrifugation method was performed in parallel to compare the synaptic protein enrichment levels. The conventional Percoll^®^ F3 and F4 synaptosome enriched fractions are shown. **p < 0.001; ***p < 0.0001. **B)** Heatmaps of the normalized abundances of the same 30 synaptic markers as used in A for each fraction and tissue or enrichment protocol. P1: pellet from the first centrifugation, containing mainly whole cells, cell debris and nuclei to serve as a negative control. **C)** Western blots of the synaptic marker SYP in differential centrifugation fractions from human forebrain organoids and human cortical tissue, at same scale to enable direct comparison between levels in fractions from FBOs and human cortex.

Consistent enrichment of synaptic proteins was observed in all pellets (P5,000, P12,000 and P22,000) from all tissue types. The highest enrichment was observed in P12,000 and P22,000 fractions from organoids and newborn mouse brain, with fold changes (35-65% enrichment) comparable to Percoll-based synaptosome fractions (40-50% enrichment, see **Supplementary table S2** for details). P22,000 fractions generally showed a higher variation. The adult brain samples (human and mouse W8) showed weaker or no enrichment in the P22,000 fractions, likely due to developmental differences in synaptic structure and composition affecting synaptosome density. Synaptic protein levels in the cytosolic fractions were depleted, confirming successful pelleting of synaptic compartments.

A heatmap of individual protein abundance (**fig. 3B**) further highlighted a trend of higher postsynaptic protein enrichment in P22,000 fractions from organoids and newborn mouse (P2), while presynaptic proteins were more prominent in P5,000 and P12,000 fractions. E.g., the presynaptic proteins Protein piccolo (PCLO), Bassoon (BSN) and SYN1 showed significant enrichment of 2.3, 1.7, and 1.5 FC respectively in the ALI-CO day 150 P12,000 fraction, while postsynaptic proteins like Glutamate receptor 1 (GRIA1) and Metabotropic glutamate receptor 2 (GRM2) showed significant enrichment of 1.5 and 1.7 FC respectively in the P22,000 fraction). This distribution pattern of pre- and postsynaptic proteins was not conserved in fractions from human cortex and adult mouse brain (**fig. 3B** and **Supplementary S3B**), possibly reflecting developmental differences between the young and adult stages, or the composition and density of the applied tissues (whole brain vs organoids).

Western blot analysis of the presynaptic marker SYP supported the mass spectrometry data, showing increased levels in P5,000 and P12,000 fractions from both FBOs and human cortex (**fig. 3C**). As expected, the more mature human cortex samples exhibited higher SYP levels than FBOs.

### Enrichment of growth cone markers

Given the early developmental stage of NOs, they are expected to contain immature synapses and actively growing axons tipped by growth cones that undergo structural and molecular changes that lead to the establishment of functional synaptic connections when they reach their final target. To assess whether the differential centrifugation protocol also enriches growth cone structures, we analyzed the distribution of 28 growth cone marker proteins (**Supplementary table S3**), curated from two proteomic studies (Estrada-Bernal et al., 2012; Nozumi et al., 2009).

In young tissue types (mouse P2, FBOs, ALI-COs day 90 and 150), growth cone proteins were significantly enriched in all pellet fractions (P5,000, P12,000, and P22,000) and depleted in the cytosolic fractions (**fig. 4A**, see details in **Supplementary table S4**). The highest overall enrichment was observed in the P12,000 fractions of organoids (41–47% increase), while in mouse P2, P22,000 showed higher average enrichment (42% increase) than P12,000 (26% increase), however with greater variability.

**Figure 4:**
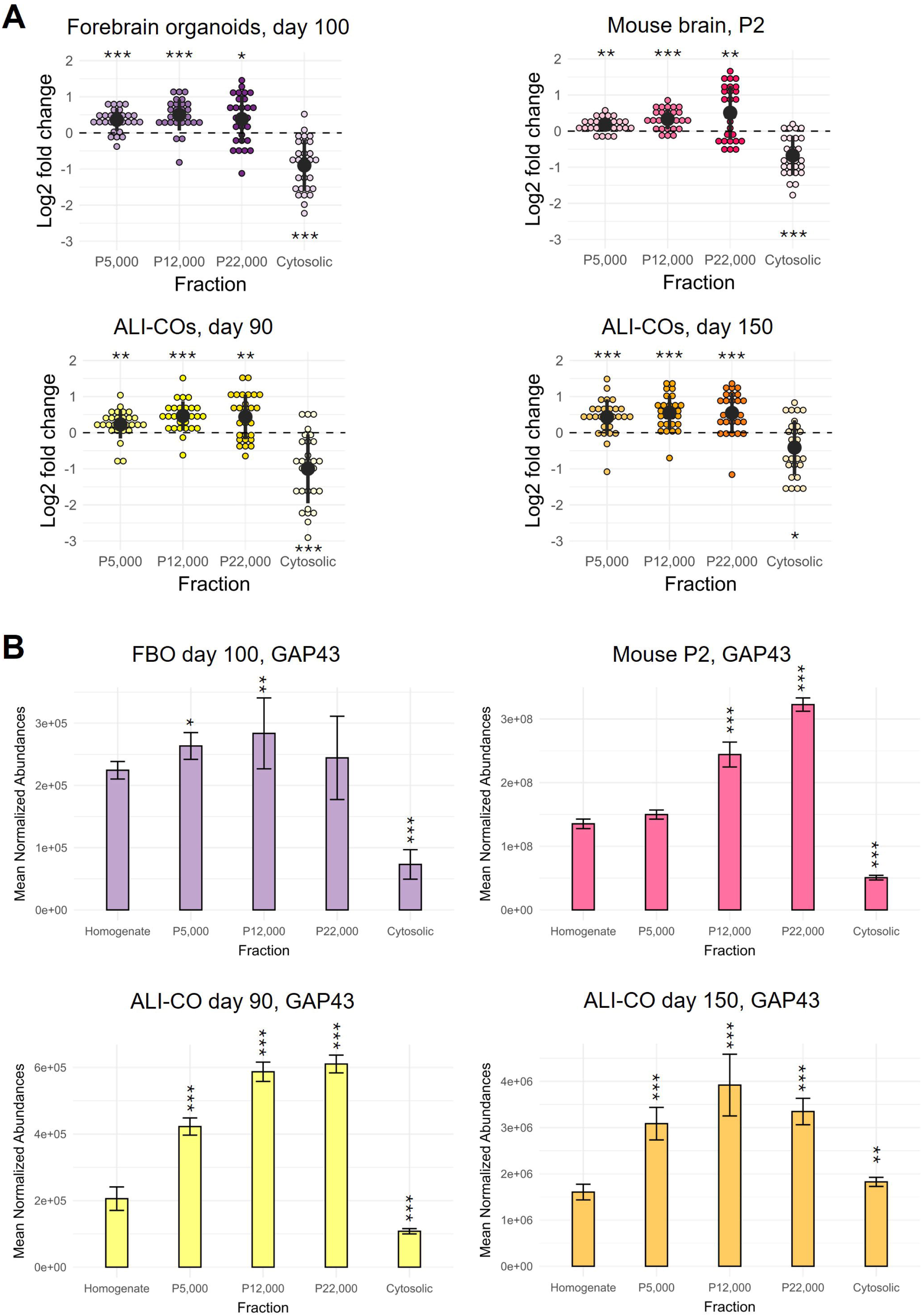
**A)** Levels of selected growth cone markers in the fractions compared to homogenate (organoids and newborn mouse (P2)). The proteins were selected based on two other proteomic studies determining markers of growth cones. *p<0.01, **p < 0.001 and ***p < 0.0001. **B)** Abundance levels of the well-known growth cone marker GAP43 in the homogenate and the four fractions derived from ALI-CO day 90 and 150, FBO and newborn mouse brain (P2). Stars in **B** indicate significant FC compared to the corresponding Homogenate level with *p < 0.05, **p < 0.01 and ***p < 0.0001.

Focusing on the canonical growth cone marker Neuromodulin (GAP43), we observed clear enrichment in the P12,000 fraction across all young tissues, with peak levels in the P22,000 fractions from ALI-CO day 90 and mouse P2 (**fig. 4B**). E.g. GAP43 showed a significant FC of 2.8 in the P12,000 fraction from day 90 ALI-COs and 2.5 from day 150 ALI-COs. These findings indicate that the workflow not only enriches synaptic material but also captures developing growth cones, the precursor of synaptosomes.

### TEM evaluation of synaptosome structures

Apart from functional GO terms and protein enrichments, the morphology of the fractions was next investigated. Functional synaptosomes require intact, membrane-enclosed structures capable of maintaining a membrane potential, depolarization and vesicle release. To verify the structural integrity of the enriched particles, we examined the P5,000, P12,000, and P22,000 fractions from human cortex, adult mouse brain (W8), and ALI-COs using transmission electron microscopy (TEM).

In adult brain samples (human and mouse), synaptosomes with clearly identifiable mitochondria and synaptic vesicles were readily observed in both the P5,000 and P12,000 fractions (human cortex in **fig. 5A-B**; mouse W8 and wider view of the same fields in **Supplementary fig. S4A-B and S4D-E**). Some synaptosomes also showed electron-dense active zones (AZs) with attached postsynaptic densities (PSDs) (**Supplementary fig. S4A and S4D**).

**Figure 5:**
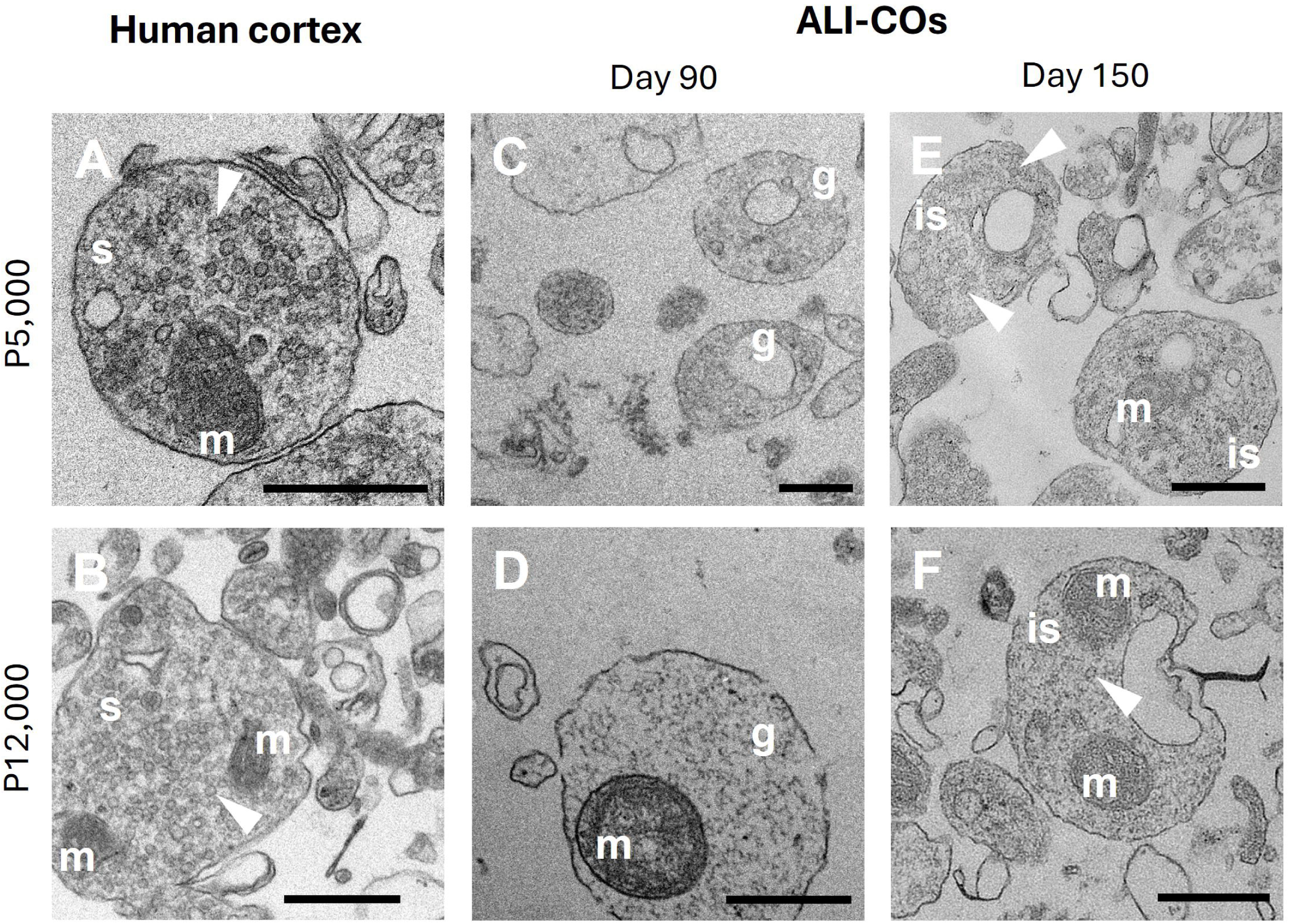
Transmission electron microscopy (TEM) images of differential centrifugation fractions from human cortex and ALI-COs at day 90 and 150. P5,000 and P12,000 fractions from human cortex **(A-B)** showing mature synaptosomes, and from ALI-COs **(C-F)** showing the presence of growth cone particles (GCPs) at day 90 and immature synaptosomes at day 150. Scalebars: 500nm. is: immature synaptosome, s: synaptosome, m: mitochondria, g: GCP, white arrows: synaptic vesicles.

ALI-COs at day 90 showed a few small synaptosome-like structures in the P5,000 and P12,000 fractions together with larger membrane-enclosed structures presenting with fibrous content, large vesicles, and mitochondria (**fig. 5C-D**, **Supplementary S4F-G**), consistent with previously described growth cone particles (GCPs) (Gordon-Weeks & Lockerbie, 1984; Grove et al., 1973; Pfenninger et al., 1983). This interpretation is supported by the observed enrichment of growth cone marker proteins, including GAP43 (**fig. 4A-B**).

By day 150, ALI-CO P5,000 and P12,000 fractions contained intact synaptosomes with mitochondria and synaptic vesicles (**fig. 5E-F**). These vesicles were fewer in number compared to those in mature synapses, aligning with the known scarcity of synaptic vesicles during early synaptogenesis (Mozhayeva et al., 2002). Accordingly, we refer to these structures as immature synaptosomes.

Both the P5,000 and the P12,000 fractions contained non-synaptic mitochondria (e.g., **Supplementary fig. S4A, D and F**). Finally, the P22,000 fractions (**Supplementary fig. S4C and J**) predominantly contained smaller membrane-bound particles, in line with the proteomic GO Cellular Component enrichment analysis showing a large representation of endomembrane and general membrane-associated terms in P22,000 fractions.

### Phosphoproteomic response to KCl depolarization of immature synaptic structures from ALI-COs

If synaptosomes are sealed and maintain a membrane potential then they can be considered metabolically active, such that KCl stimulation will result in membrane depolarization and subsequent release of neurotransmitters by Ca^2+^ mediated exocytosis followed by ultra-fast endocytosis of the synaptic vesicles (Silbern et al., 2021). These molecular functions are tightly controlled by protein phosphorylation and dephosphorylation via multiple kinases and phosphatases (Engholm-Keller et al., 2019; Kohansal-Nodehi et al., 2016; Silbern et al., 2021). To test whether the derived GCPs and immature synaptosomes were metabolically active and capable of stimulation-induced exo-/endocytosis, we stimulated the P5,000 and P12,000 fractions of the NOs, using a high concentration of KCl (76.2 mM). Stimulation of synaptic structures derived from the FBOs did not yield any significant changes in phosphorylation, likely due to too low amount of material (Data not shown). ALI-COs generally yielded more material per tissue gram, though this tendency was not statistically significant (**Supplementary fig. S4K-L**). Stimulation of enriched synaptic structures from the ALI-COs at day 90 and 150 resulted in significant regulation of phosphorylation in 219 proteins at day 90, and 73 proteins at day 150 with an overlap of 16 proteins comprising proteins important for exo-/endocytosis at the synapse, like Dynamin1 (DNM1), and PCLO, cytoskeletal proteins Doublecortin (DCX), Microtubule-Associated Protein 1B (MAP1B), and the synaptic scaffold protein Disks large-associated protein 4 (DLGAP4) (**fig. 6A**). A well-described, key indication of active endocytosis in synaptosomes is the dephosphorylation of S744/S778 phosphosites in DNM1, which allows scission and release of endocytosed vesicles from the membrane (Cousin & Robinson, 2001; Graham et al., 2007). At both timepoints, DNM1 shows significant dephosphorylation of the doubly phosphorylated S774/S778 peptide, indicating active endocytosis (**fig. 6B**).

**Figure 6:**
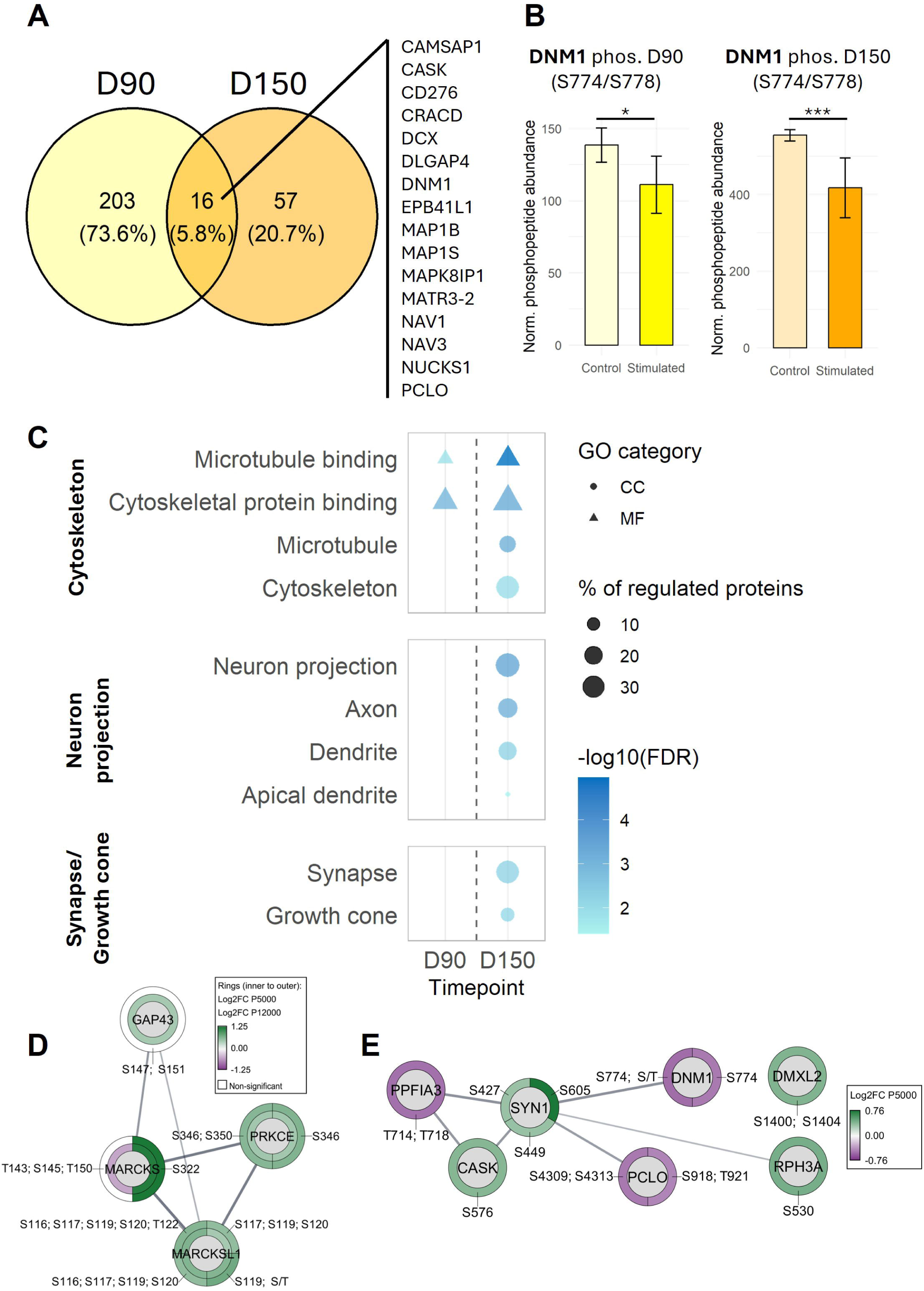
**A)** Venn diagram of proteins with significant KCl stimulation induced changes in phosphorylation (P5,000 and P12,000 fraction proteins combined) showing the overlapping proteins with regulation in both day 90 and day 150 ALI-CO fractions. **B)** DNM1 S774 or S774/S778 down-regulation was detected at both timepoints (P5,000 fraction at day 150; P12,000 fraction at day 90). **C)** GO molecular function (MF) or Cellular Compartment pathway enrichment among the proteins with stimulation-dependent phospho-regulation in ALI-CO fractions from day 90 and 150. **D)** PPI STRING network (functional associations, incl. text mining) of growth cone proteins and their changes in phosphorylation at day 90. **E)** PPI STRING network (functional associations, incl. text mining) of proteins involved in synaptic vesicle cycling (extracted using the SynGO portal online tool) and their stimulation-dependent changes in phosphorylation in day 150 ALI-CO fractions.

GO enrichment analysis of proteins with stimulation-dependent changes in phosphorylation showed a difference in stimulation profiles of day 90 and 150. Stimulation at day 90 primarily showed changes within cytoskeleton terms, while at day 150, the proteins with phospho-regulations were enriched in both cytoskeleton, neuron projection, and synapse/growth cone terms (**fig. 6C**). Characteristic for the regulation of phosphorylation in the day 90 fractions, was a cluster of proteins specific for growth cones and important for growth cone motility (MARCKS (Myristoylated alanine-rich C-kinase substrate), MARCKSL1 (MARCKS-related protein), PRKCE (Protein kinase C epsilon type) and GAP43, **fig. 6D**) (Gatlin et al., 2006; Nozumi et al., 2009). GAP43 also plays an important role in Ca^2+^ signaling at the synapse (Haruta et al., 1997). Furthermore, proteins involved in neurite outgrowth and axonal elongation showed changes in phosphorylation in day 90 fractions including DCLK1 (Serine/threonine-protein kinase DCLK1), SHTN1 (Shootin 1), MARK4 (MAP/microtubule affinity-regulating kinase 4) and MAP1B. In synaptic structures from day 150 ALI-FOs, consistent with the later developmental stage and presence of synaptic vesicles in the enriched structures, we observed stimulation-dependent phosphorylation changes in proteins involved in the synaptic vesicle cycle including exo-/endocytosis proteins (e.g., PCLO, DNM1, SYN1, RPH3A (Rabphilin 3A) and CASK (Peripheral plasma membrane protein CASK), **fig. 6E**). Other proteins showing changes in phosphorylation were MAPT (Microtubule-associated protein tau), CAMK4 (Calcium/calmodulin-dependent protein kinase type IV) and DLGAP4 which all play a role in synaptic signaling and/or plasticity (Guo et al., 2017; Ho et al., 2000; Pavinato et al., 2023; Won et al., 2017) (Rasmussen et al., 2017). .

Among the proteins with regulations in phosphorylation at both timepoints are also proteins associated with neurodevelopmental and psychiatric disorders such as autism spectrum disorders (SHANK1 (SH3 and multiple ankyrin repeat domains protein 1) (Gong & Wang, 2015), DLGAP4 (Schob et al., 2019) and MAP1B (Liu et al., 2015)), intellectual disability (MAP1B (Liu et al., 2015; Walters et al., 2018), CAMK2G (Calcium/calmodulin-dependent protein kinase type II subunit gamma) (Proietti Onori et al., 2018), DLGAP4 (Schob et al., 2019) and CASK (Moog & Kutsche, 1993)) and Schizophrenia (DCLK1 (Håvik et al., 2012), MARCKS (Pinner et al., 2014), PRKCE (Zhao et al., 2013) and SHANK1 (Fromer et al., 2014)).

## Discussion

Our study establishes a differential centrifugation protocol that enables isolation of functional GCPs and immature synaptosomes from human NOs, addressing critical limitations of traditional synaptosome enrichment methods for this kind of material. By integrating proteomic, ultrastructural, and functional analyses, we demonstrate that ALI-COs recapitulate key stages of synaptic development: day 90 fractions showing stimulation-dependent changes in cytoskeletal regulators (e.g., MARCKS, MARCKSL1) and growth cone markers (GAP43), while day 150 fractions transition toward immature synaptosomes with changes in phosphorylation in synaptic vesicle cycle proteins (e.g., SYN1, RPH3A) and the key ultra-fast endocytosis protein DNM1. This temporal progression is consistent with the expected features of *in vivo* synaptogenesis, where axonal growth and pathfinding precedes synaptic maturation, providing a human-specific model to study synaptic development and dysfunction.

### Methodological Advancements and Validation

The simplicity of the protocol, bypassing density gradients, and the need for only minimal tissue volume allows scalability across species and organoid models. Vesicle-dense synaptosomes were successfully enriched from human cortical tissue in the P5,000 and P12,000 fractions as confirmed by TEM, demonstrating the effectiveness of the method even with limited tissue samples and relatively immature organoids.

Proteomic profiling revealed robust enrichment of synaptic proteins in P5,000 and P12,000 fractions (e.g., 2.3-fold increase in PCLO in day 150 ALI-COs vs. homogenate), while non-synaptic mitochondrial contamination was predominantly confined to P5,000 fractions (validated with TOMM20). The P22,000 fractions showed enrichment in postsynaptic proteins and endomembrane structures. Ultrastructural analysis by TEM confirmed membrane-enclosed GCPs with fibrous content resembling actin networks, larger vesicles and mitochondria in day 90 isolates, which is consistent with other other studies on GCPs (Gordon-Weeks & Lockerbie, 1984; Grove et al., 1973; Pfenninger et al., 1983). Day 150 fractions contained immature synaptosomes marked by sparse synaptic vesicles. These findings align with the expected developmental shifts in NOs over time.

### Functional Maturation and Phosphoproteomic Signatures

Functional viability was validated through KCl-induced depolarization of synaptic structures isolated from ALI-COs. This triggered phosphorylation changes in growth cone markers (e.g. GAP43, MARCKS, MARCKSL1), cytoskeletal regulators (DCLK1, SHTN1, MARK4), and calcium-dependent kinases (CAMK2G, PRKCE) in day 90 GCPs. These phosphorylation dynamics align with established mechanisms of growth cone motility, in which Ca²⁺ influx modulates actin-microtubule interactions (Henley & Poo, 2004; Nelson et al., 2013; Schneider et al., 2023). Rodent studies have demonstrated that growth cones and isolated GCPs can depolarize in response to stimulation, leading to Ca²⁺ transients and cytoskeleton remodeling, which is highly controlled by phosphorylation (Belardetti et al., 1986; Lipscombe et al., 1988; Lockerbie et al., 1991; Neely & Gesemann, 1994). It is e.g. known that PRKCE-dependent MARCKS phosphorylation is involved in growth cone adhesion and turning (Gatlin et al., 2006). This supports our interpretation that the day 90 phosphorylation profile reflects a growth cone–dominated stimulation response. In day 150 fractions, depolarization induced regulation of phosphorylation in synaptic vesicle cycling-related proteins (SYN1, DNM1, CASK, RPH3A, DMXL2). GO enrichment analysis showed enrichment in cytoskeleton, synaptic and neuron projection related terms, indicating a more mature stimulation profile compared to day 90 fractions. Importantly, DNM1 S774/S778 dephosphorylation, a strong indicator of active endocytosis and synaptic vesicle cycling (Cousin & Robinson, 2001; Graham et al., 2007), was observed in both day 90 and day 150 fractions.

The functional changes in protein phosphorylation in KCl stimulated synaptic structures from ALI-COs, which was not possible to show in FBO isolates, could indicate a higher level of maturity in the ALI cultures, which have previously been shown (Giandomenico et al., 2021). Higher number of synapses/growth cones and/or more mature, spine-like morphology could result in a better quality of the enriched structures. The lack of significant changes in phosphorylation upon KCl stimulation of structures from FBOs could however also be due to too low amount of material.

### Implications for Disease Modeling and Functional Studies

The protocol offers value beyond structural enrichment. Phosphoproteomic profiling of day 150 ALI-COs identified phosphorylation changes in neurodevelopmental disorder-associated proteins (e.g., SHANK1, DLGAP4 and CAMK2G), offering a platform to study synaptic dysfunction in conditions like autism spectrum disorder, intellectual disability and schizophrenia. For instance, CAMK2G is a protein kinase implicated in intellectual disability (Proietti Onori et al., 2018), which here showed depolarization-responsive phosphorylation, suggesting its role in activity-dependent synaptic maturation. Similarly, the stimulation-dependent regulation of phosphorylation in MARCKS and MARCKSL1, cytoskeletal modulators linked to neuronal migration defects (El Amri et al., 2018), underscores the method’s potential to probe early pathogenic mechanisms.

The method’s compatibility with patient-derived organoids enables direct investigation of disease-relevant synaptic phenotypes in a human context. In parallel, CRISPR/CAS approaches can be used to model specific gene perturbations or SNPs (e.g. in *SHANK3* or *NRXN1*) allowing systematic dissection of synaptic gene function. Furthermore, the ability to isolate metabolically active GCPs permits investigations into axon guidance responses (e.g., semaphorin and netrin signaling), bypassing ethical and logistical constraints of postmortem or primary human tissue.

### Limitations and Future Directions

In addition to developmental stage differences, a key distinction between brain organoids and native brain tissue lies in their structural and cellular composition. Mature brain tissue is densely packed and includes abundant white matter, comprising myelinated axons and oligodendrocytes, as well as astrocytes, microglia, and vasculature. These all contribute to tissue density, biochemical complexity, and centrifugation behavior. Most neural organoids, in contrast, lack myelination and supporting glial populations, particularly oligodendrocytes and mature astrocytes, which emerge only under prolonged culture or directed differentiation protocols (Paşca et al., 2015; Madhavan et al., 2018). The absence of white matter tracts and compact axon bundles likely reduces physical resistance during homogenization and may shift the sedimentation profiles of synaptic components, particularly in differential centrifugation protocols. The reduced extracellular matrix and lower tissue stiffness of organoids compared to intact brain (Lancaster et al., 2013; Quadrato et al., 2017) may lead to more efficient extraction of membrane-bound structures but may also influence organelle integrity. These structural differences should be considered when comparing subcellular enrichment yields and interpreting the spatial distribution of synaptic and cytoskeletal proteins.

The relatively limited number of depolarization-responsive phosphorylation events correlates with the low presence of GCPs and immature synaptosomes detected by electron microscopy, suggesting that only a small fraction of physiologically sealed, metabolically active synaptosomes are present. This may reflect limited structural maturity, suboptimal purification, or biophysical differences in lipid or protein composition between NOs and native brain tissue. While extra-synaptosomal mitochondrial contamination in P5,000 fractions remains a challenge, the P12,000 fraction provides the purest enrichment of functional synaptic structures for stimulation experiments. Further optimization of fractionation protocols or incorporation of immunoaffinity-based purification strategies could help to further reduce non-synaptic contaminants.

Additionally, extending organoid culture beyond day 150 may yield more synaptosomes with mature synaptic vesicle pools, resulting in higher yield of functional active synaptosomes and thereby enabling studies of neurotransmitter release kinetics. Systematic evaluation of batch-to-batch organoid variability, regional patterning, and synapse subtype composition may be critical to increase reproducibility and translational relevance. Furthermore, variable yield of enriched synaptic structures between organoid models (e.g. spheric vs. ALI cultures) will affect the scalability of the method. These will be important aspects to address for effective scalability of the model for e.g. drug screening. The protocol’s reproducibility across human and murine tissues underscores its versatility. Coupled with advances in spatial proteomics, this method could be deployed to map subcellular proteome dynamics during synaptic maturation or drug responses. For example, integration of synaptosome depolarization with single-organoid real-time calcium imaging could offer deeper insight into human-specific signaling cascades underlying synaptic plasticity. Since this model enables access to early-stage synaptic structures, it may also allow future investigation into upstream regulators of synaptogenesis, such as guidance cues, adhesion molecules, and cytoskeletal remodeling enzymes.

## Conclusion

By integrating a differential centrifugation strategy with functional and proteomic profiling, we have established a practical framework for isolating active synaptic structures from human neural organoids. Th approach reveals temporally distinct molecular signatures that reflect developmental transitions in synapse formation and enables the identification of phosphorylation events linked to growth cone remodeling and synaptic vesicle cycling. Beyond enriching specific compartments, the protocol facilitates future mechanistic interrogation of synaptic signaling, with the potential to be adapted for high-throughput studies and for patient-specific disease models. As cerebral organoid technology matures, this platform is anticipated to provide a new opportunity to explore human synaptic development and dysfunction in an experimentally accessible, physiologically responsive system.

## Acknowledgements

The project was funded by the Independent Research Fund Denmark in the category of Natural Science (project no. 9040-00381B) and by the Lundbeck Foundation under the DEVELOPNOID project (project no. R336+2020-1113). Immunofluorescent image acquisition was conducted at the Danish Molecular Biomedical Imaging Center (DaMBIC, University of Southern Denmark), supported by the Novo Nordisk Foundation (NNF) (grant agreement no. NNF18SA0032928). This project was supported by a generous grant from the Danish Agency of Higher Education and Science to establish the PLATO research infrastructure: Danish National Mass Spectrometry Platform for Proteomics and Biomolecular Imaging (grant no. 5229-00012B, www.sdu.dk/PLATO). PJR was supported by the National Health & Medical Research Council Australia (GNT1069493, GNT1052494, and GNT1047070) and the Children’s Medical Research Institute (CMRI).

## Author contributions

MRL, MSØ, PJ, PJR, and LC designed the study and developed the method. MSØ, LC, PJ, DJLDS, MS, LAJ, MP, HB, SB and MT conducted the experiments. JB, FRP, KF and MAL provided external resources and material. MSØ, MRL, PJ and VS conducted data analysis and interpretation of the results. MSØ and PJR drafted the first manuscript supported by LC, PJ, DJLDS and MP. PJR co-supervised the project. MRL conceived the study and acquired funding for the project. All authors read, reviewed and approved the final version of the manuscript.

## Abbreviations

2D: two-dimensional
3D: three-dimensional
ACN: acetonitrile
AGC: automatic gain control
ALI: air-liquid interface
ALI-CO: air-liquid interface cerebral organoid
AZ: active zone
BP: Biological Process
BSN: Bassoon
CAMK2G: Calcium/calmodulin-dependent protein kinase II gamma
CAMK4: Calcium/calmodulin-dependent protein kinase type IV
CASK: Calcium/calmodulin-dependent serine protein kinase
CC: Cellular component
CO: cerebral organoid
DCLK1: Doublecortin-like kinase 1
DCX: Doublecortin
DDA: data dependent acquisition
DIA: data independent acquisition
DLGAP4: Disks large-associated protein 4
DNM1: Dynamin-1
DTT: dithiothreitol
EBs: embryoid bodies
ESI: electrospray ionisation
EtOH: ethanol
FA: formic acid
FBO: dorsal forebrain organoid
FC: fold change
FDR: false discovery rate
FWHM: full width half maximum
GAP43: Neuromodulin, alternative name: Growth associated protein 43
GCP: growth cone particle
GO: Gene Ontology
GRIA1: Glutamate receptor 1
GRM2: Metabotropic glutamate receptor 2
HBK: HEPES-buffered Krebs-like
HCD: higher energy collision dissociation
HPLC: high performance liquid chromatography
IAA: iodoacetamide
IHC: immunohistochemical
iPSC: induced pluripotent stem cell
LMB: Laboratory of Molecular Biology
m/z: mass to charge ratio
MAP1B: Microtubule-associated protein 1B
MAP2: Microtubule-associated protein 2
MAPT: Microtubule-associated protein tau
MARCKS: Myristoylated alanine-rich C-kinase substrate
MARCKSL1: MARCKS-related protein
MARK4: Microtubule-associated regulatory kinase 4
MF: Molecular Function
MRC: Medical Research Council
MS: mass spectrometry
MS/MS: Tandem mass spectrometry
NCE: normalized collision energy
nLC: Nano Liquid Chromatography
NO: neural organoid
ON: overnight
P1: pellet after initial 800xg centrifugation
P12,000: pellet after 20 min 12,000xg centrifugation
P2: postnatal day 2
P22,000: pellet after 20 min 22,000xg centrifugation
P5,000: pellet after 10 min 5,000xg centrifugation
PASEF: parallel accumulation serial fragmentation
PBS: phosphate buffered saline
PCA: principal component analysis
PCLO: Piccolo
PD: Proteome Discoverer
PPI: Protein-protein interaction
PRKCE: Protein kinase C type epsilon
PSC: pluripotent stem cell
PSD: postsynaptic density
PSD95: Postsynaptic density protein 95, alternative DLG4
PTM: post-translational modification
RP: reversed phase
RPH3A: Rabphilin 3A
RT: room temperature
S1: supernatant after initial 800xg centrifugation
SFSCM: serum free slice culture medium
SHANK1: SH3 and multiple ankyrin repeat domains protein 1
SHTN1: Shootin1
SP: sodium phosphate
SPS-MS3: Synchronous Precursor Selection Multi-Stage Mass Spectrometry
SYN1: Synapsin-1
SYP: Synaptophysin
TEAB: triethylammonium bicarbonate
TEM: transmission electron microscopy
TFA: trifluoracetic acid
TMBST: TBS-Tween20
TMT: Tandem Mass Tags
TOMM20: Translocase of outer mitochondrial membrane 20
VGLUT1: Vesicular glutamate transporter 1
W8: 8-week-old

