## Supplementary for "Modeling Synaptic Maturation from Growth Cone to Synapse in Human Organoids"

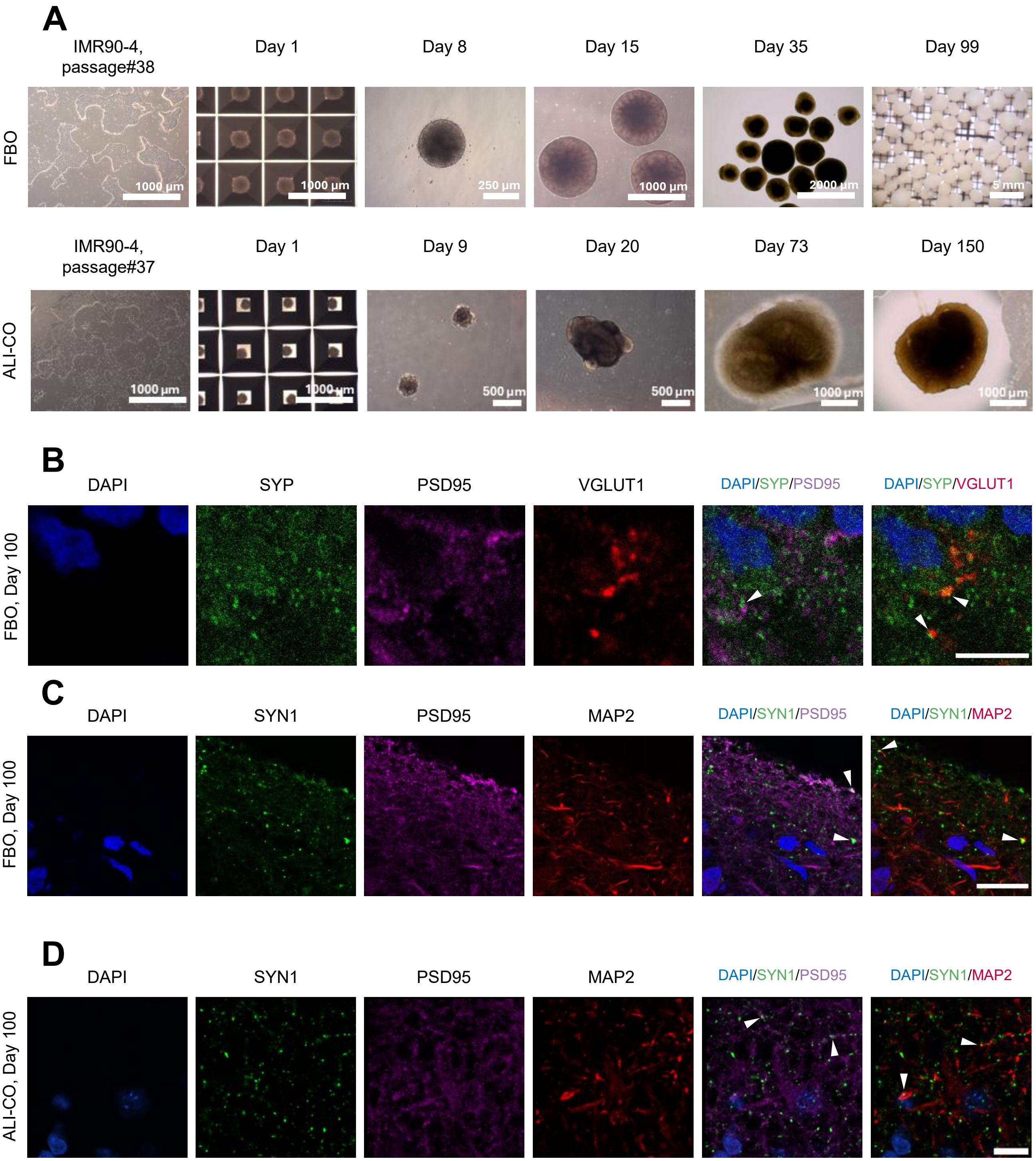


**Figure S1: A)** Light microscopic images of IMR90-4 iPSCs and forebrain organoids (FBOs) or ALI-COs at different differentiation stages. **B-C)** Immunohistochemical (IHC) labelling of synapses in FBOs (B-C) at day 100 and ALI-COs (D) at day 100. **B)** FBO day 100 labelling of presynaptic markers Synaptophysin (SYP) and Vesicular glutamate transporter 1 (VGLUT1) and postsynaptic marker Postsynaptic density protein 95 (PSD95) at day 99. Scalebar: 5µm. **C)** FBO day 100 labelling of presynaptic marker Synapsin 1 (SYN1), postsynaptic marker PSD95 and dendritic marker Microtubule associated protein 2 (MAP2) at day 100. Scalebar: 20µm. **D)** ALI-COs labeled at day 100 for same markers as in **C** (scale bar: 10µm). White arrows pointing at areas with co-localization or adjacent expression of the pre- and postsynaptic markers. For all IHC images, n=3.

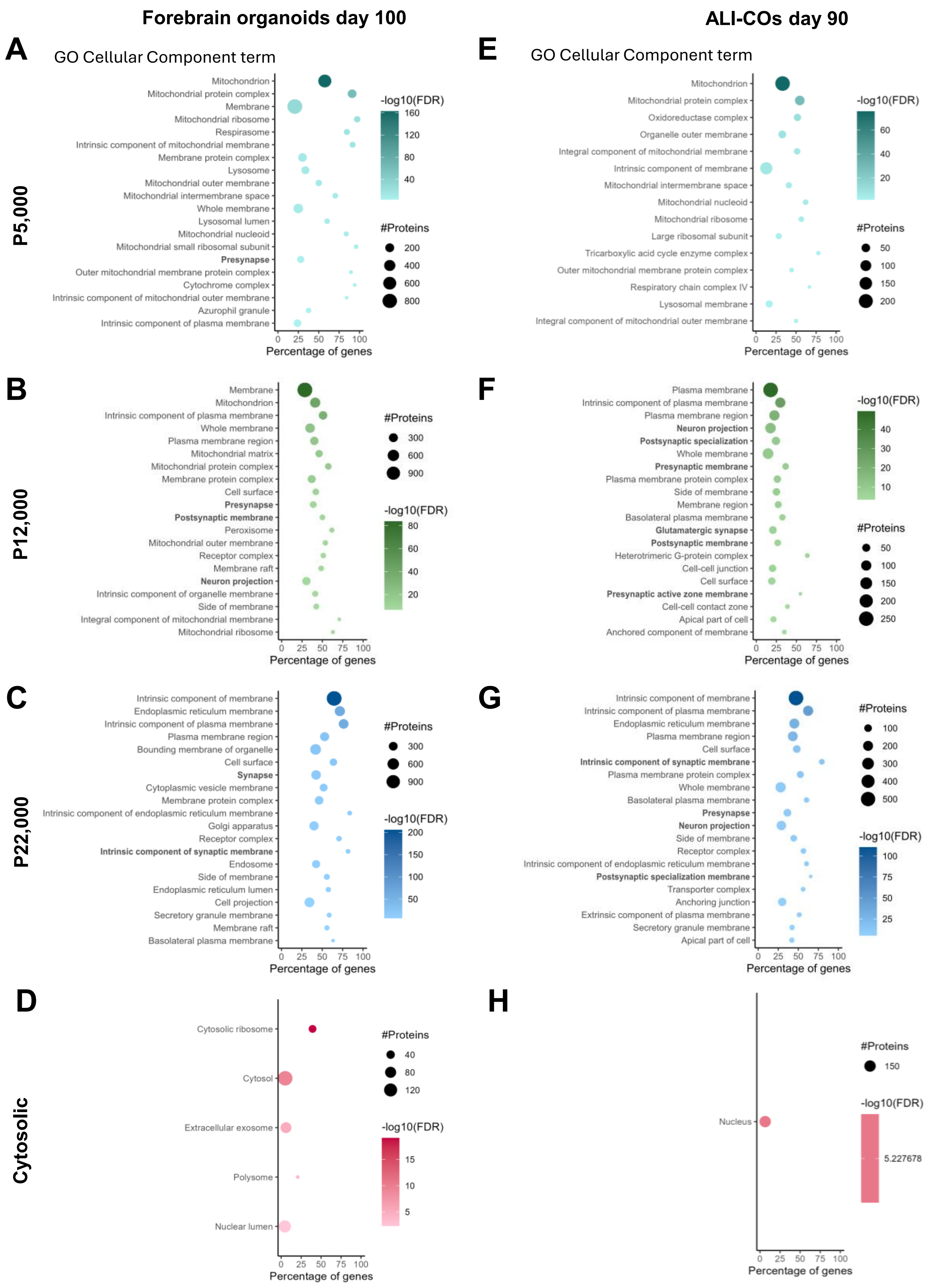

Forebrain organoids day 100
ALI-COs day 90
A
GO Cellular Component term
P5,000
Mitochondrion
Mitochondrial protein complex
Membrane
Mitochondrial ribosome
Respirasome
Intrinsic component of mitochondrial membrane
Membrane protein complex
Lysosome
Mitochondrial outer membrane
Mitochondrial intermembrane space
Whole membrane
Lysosomal lumen
Mitochondrial nucleoid
Mitochondrial small ribosomal subunit
Presynapse
Outer mitochondrial membrane protein complex
Cytochrome complex
Intrinsic component of mitochondrial outer membrane
Azurophil granule
Intrinsic component of plasma membrane
Percentage of genes
-log10(FDR)
#Proteins
E
GO Cellular Component term
Mitochondrion
Mitochondrial protein complex
Oxidoreductase complex
Organelle outer membrane
Integral component of mitochondrial membrane
Intrinsic component of membrane
Mitochondrial intermembrane space
Mitochondrial nucleoid
Mitochondrial ribosome
Large ribosomal subunit
Tricarboxylic acid cycle enzyme complex
Outer mitochondrial membrane protein complex
Respiratory chain complex IV
Lysosomal membrane
Integral component of mitochondrial outer membrane
Percentage of genes
B
P12,000
Membrane
Mitochondrion
Intrinsic component of plasma membrane
Whole membrane
Plasma membrane region
Mitochondrial matrix
Mitochondrial protein complex
Membrane protein complex
Cell surface
Presynapse
Postsynaptic membrane
Peroxisome
Mitochondrial outer membrane
Receptor complex
Membrane raft
Neuron projection
Intrinsic component of organelle membrane
Side of membrane
Integral component of mitochondrial membrane
Mitochondrial ribosome
Percentage of genes
F
Plasma membrane
Intrinsic component of plasma membrane
Plasma membrane region
Neuron projection
Postsynaptic specialization
Whole membrane
Presynaptic membrane
Plasma membrane protein complex
Side of membrane
Membrane region
Basolateral plasma membrane
Glutamatergic synapse
Postsynaptic membrane
Heterotrimeric G-protein complex
Cell-cell junction
Cell surface
Presynaptic active zone membrane
Cell-cell contact zone
Apical part of cell
Anchored component of membrane
Percentage of genes
C
P22,000
Intrinsic component of membrane
Endoplasmic reticulum membrane
Intrinsic component of plasma membrane
Plasma membrane region
Bounding membrane of organelle
Cell surface
Synapse
Cytoplasmic vesicle membrane
Membrane protein complex
Intrinsic component of endoplasmic reticulum membrane
Golgi apparatus
Receptor complex
Intrinsic component of synaptic membrane
Endosome
Side of membrane
Endoplasmic reticulum lumen
Cell projection
Secretory granule membrane
Membrane raft
Basolateral plasma membrane
Percentage of genes
G
Intrinsic component of membrane
Intrinsic component of plasma membrane
Endoplasmic reticulum membrane
Plasma membrane region
Cell surface
Intrinsic component of synaptic membrane
Plasma membrane protein complex
Whole membrane
Basolateral plasma membrane
Presynapse
Neuron projection
Side of membrane
Receptor complex
Intrinsic component of endoplasmic reticulum membrane
Postsynaptic specialization membrane
Transporter complex
Anchoring junction
Extrinsic component of plasma membrane
Secretory granule membrane
Apical part of cell
Percentage of genes
D
Cytosolic
Cytosolic ribosome
Cytosol
Extracellular exosome
Polysome
Nuclear lumen
Percentage of genes
H
Nucleus
#Proteins
150
-log10(FDR)
5.227678
Percentage of genes

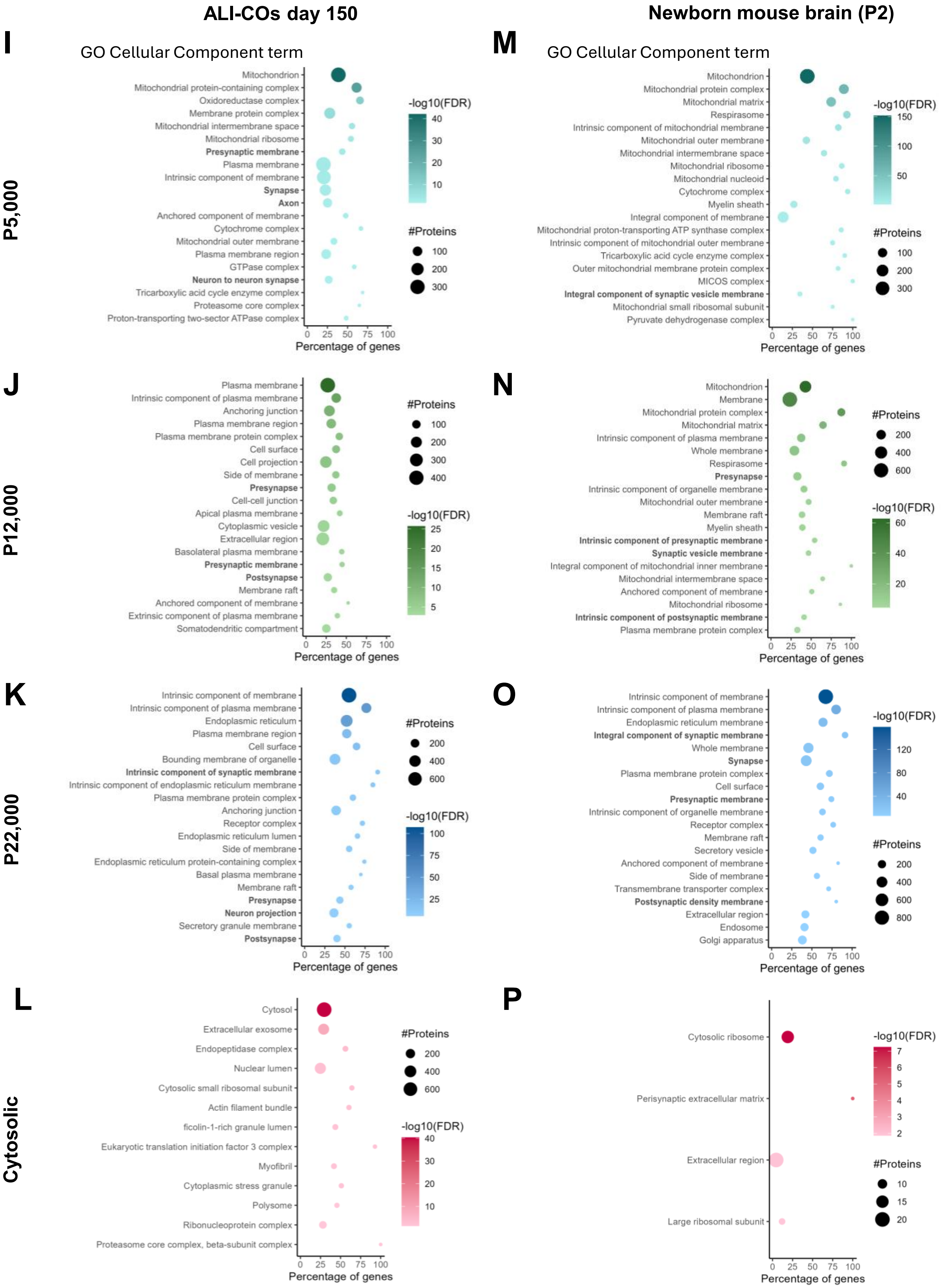

ALI-COs day 150
Newborn mouse brain (P2)
I
GO Cellular Component term
Mitochondrion
Mitochondrial protein-containing complex
Oxidoreductase complex
Membrane protein complex
Mitochondrial intermembrane space
Mitochondrial ribosome
Presynaptic membrane
Plasma membrane
Intrinsic component of membrane
Synapse
Axon
Anchored component of membrane
Cytochrome complex
Mitochondrial outer membrane
Plasma membrane region
GTPase complex
Neuron to neuron synapse
Tricarboxylic acid cycle enzyme complex
Proteasome core complex
Proton-transporting two-sector ATPase complex
-log10(FDR)
40
30
20
10
#Proteins
100
200
300
0 25 50 75 100
Percentage of genes
P5,000
M
GO Cellular Component term
Mitochondrion
Mitochondrial protein complex
Mitochondrial matrix
Respirasome
Intrinsic component of mitochondrial membrane
Mitochondrial outer membrane
Mitochondrial intermembrane space
Mitochondrial ribosome
Mitochondrial nucleoid
Cytochrome complex
Myelin sheath
Integral component of membrane
Mitochondrial proton-transporting ATP synthase complex
Intrinsic component of mitochondrial outer membrane
Tricarboxylic acid cycle enzyme complex
Outer mitochondrial membrane protein complex
MICOS complex
Integral component of synaptic vesicle membrane
Mitochondrial small ribosomal subunit
Pyruvate dehydrogenase complex
-log10(FDR)
150
100
50
#Proteins
100
200
300
0 25 50 75 100
Percentage of genes
J
Plasma membrane
Intrinsic component of plasma membrane
Anchoring junction
Plasma membrane region
Plasma membrane protein complex
Cell surface
Cell projection
Side of membrane
Presynapse
Cell-cell junction
Apical plasma membrane
Cytoplasmic vesicle
Extracellular region
Basolateral plasma membrane
Presynaptic membrane
Postsynapse
Membrane raft
Anchored component of membrane
Extrinsic component of plasma membrane
Somatodendritic compartment
#Proteins
100
200
300
400
-log10(FDR)
25
20
15
10
5
0 25 50 75 100
Percentage of genes
P12,000
N
Mitochondrion
Membrane
Mitochondrial protein complex
Mitochondrial matrix
Intrinsic component of plasma membrane
Whole membrane
Respirasome
Presynapse
Intrinsic component of organelle membrane
Mitochondrial outer membrane
Membrane raft
Myelin sheath
Intrinsic component of presynaptic membrane
Synaptic vesicle membrane
Integral component of mitochondrial inner membrane
Mitochondrial intermembrane space
Anchored component of membrane
Mitochondrial ribosome
Intrinsic component of postsynaptic membrane
Plasma membrane protein complex
#Proteins
200
400
600
-log10(FDR)
60
40
20
0 25 50 75 100
Percentage of genes
K
Intrinsic component of membrane
Intrinsic component of plasma membrane
Endoplasmic reticulum
Plasma membrane region
Cell surface
Bounding membrane of organelle
Intrinsic component of synaptic membrane
Intrinsic component of endoplasmic reticulum membrane
Plasma membrane protein complex
Anchoring junction
Receptor complex
Endoplasmic reticulum lumen
Side of membrane
Endoplasmic reticulum protein-containing complex
Basal plasma membrane
Membrane raft
Presynapse
Neuron projection
Secretory granule membrane
Postsynapse
#Proteins
200
400
600
-log10(FDR)
100
75
50
25
0 25 50 75 100
Percentage of genes
P22,000
O
Intrinsic component of membrane
Intrinsic component of plasma membrane
Endoplasmic reticulum membrane
Integral component of synaptic membrane
Whole membrane
Synapse
Plasma membrane protein complex
Cell surface
Presynaptic membrane
Intrinsic component of organelle membrane
Receptor complex
Membrane raft
Secretory vesicle
Anchored component of membrane
Side of membrane
Transmembrane transporter complex
Postsynaptic density membrane
Extracellular region
Endosome
Golgi apparatus
-log10(FDR)
120
80
40
#Proteins
200
400
600
800
0 25 50 75 100
Percentage of genes
L
Cytosol
Extracellular exosome
Endopeptidase complex
Nuclear lumen
Cytosolic small ribosomal subunit
Actin filament bundle
ficolin-1-rich granule lumen
Eukaryotic translation initiation factor 3 complex
Myofibril
Cytoplasmic stress granule
Polysome
Ribonucleoprotein complex
Proteasome core complex, beta-subunit complex
#Proteins
200
400
600
-log10(FDR)
40
30
20
10
0 25 50 75 100
Percentage of genes
Cytosolic
P
Cytosolic ribosome
Perisynaptic extracellular matrix
Extracellular region
Large ribosomal subunit
-log10(FDR)
7
6
5
4
3
2
#Proteins
10
15
20
0 25 50 75 100
Percentage of genes

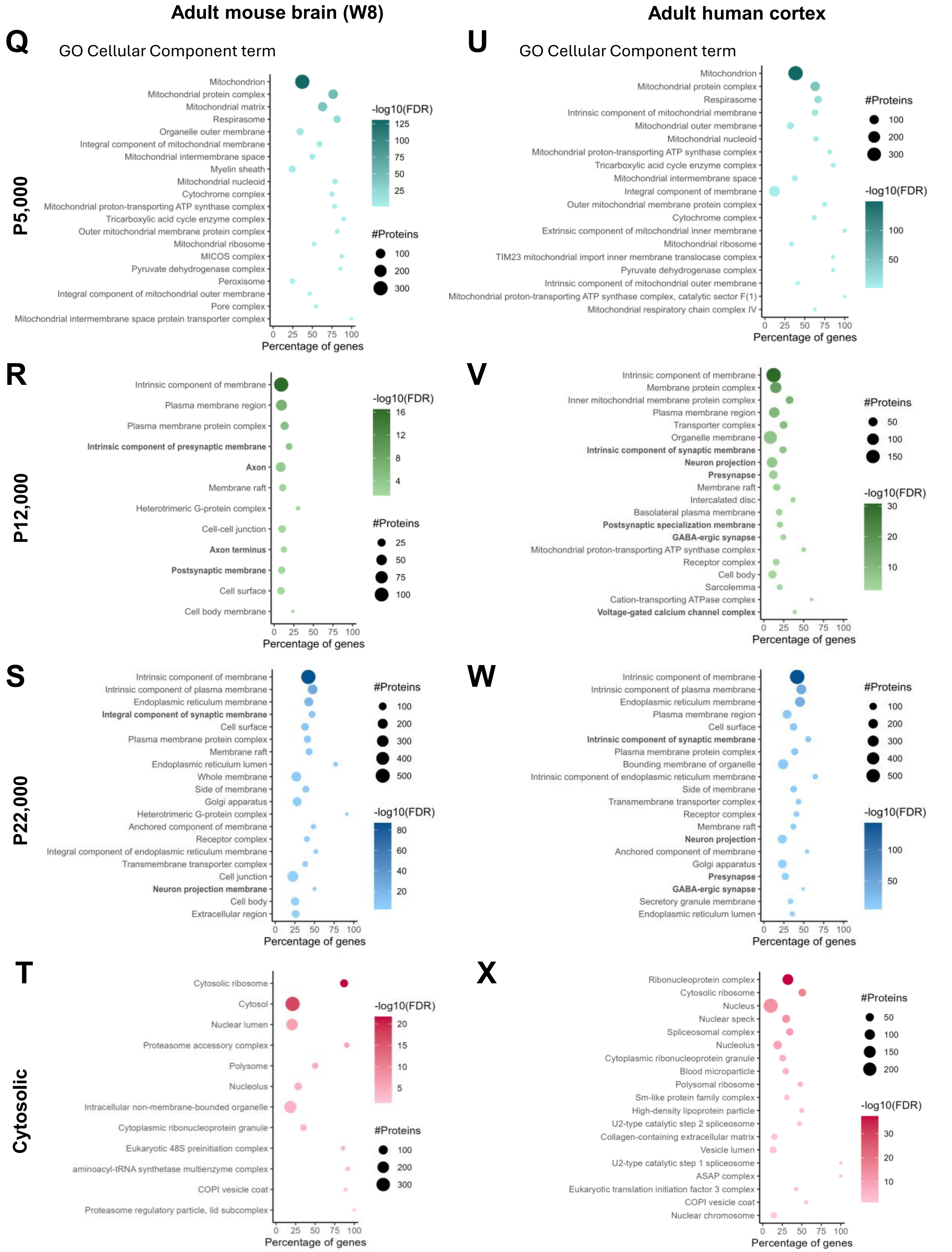


Figure S2: A-X Gene Ontology (GO) Cellular Component enriched terms (top 20, FDR ≤ 0.05) based on fraction-specific proteins (FC ≥ 1.5 compared to homogenate, FDR ≤ 0.01). P5,000, P12,000, P22,000 and Cytosolic fractions from differential centrifugation of forebrain organoids (day 100), ALI-COs day 90, ALI-COs day 150, newborn mouse brain tissue (postnatal day 2 (P2)), adult mouse brain tissue (week 8 (W8)) and surgery derived adult human cortical tissue.

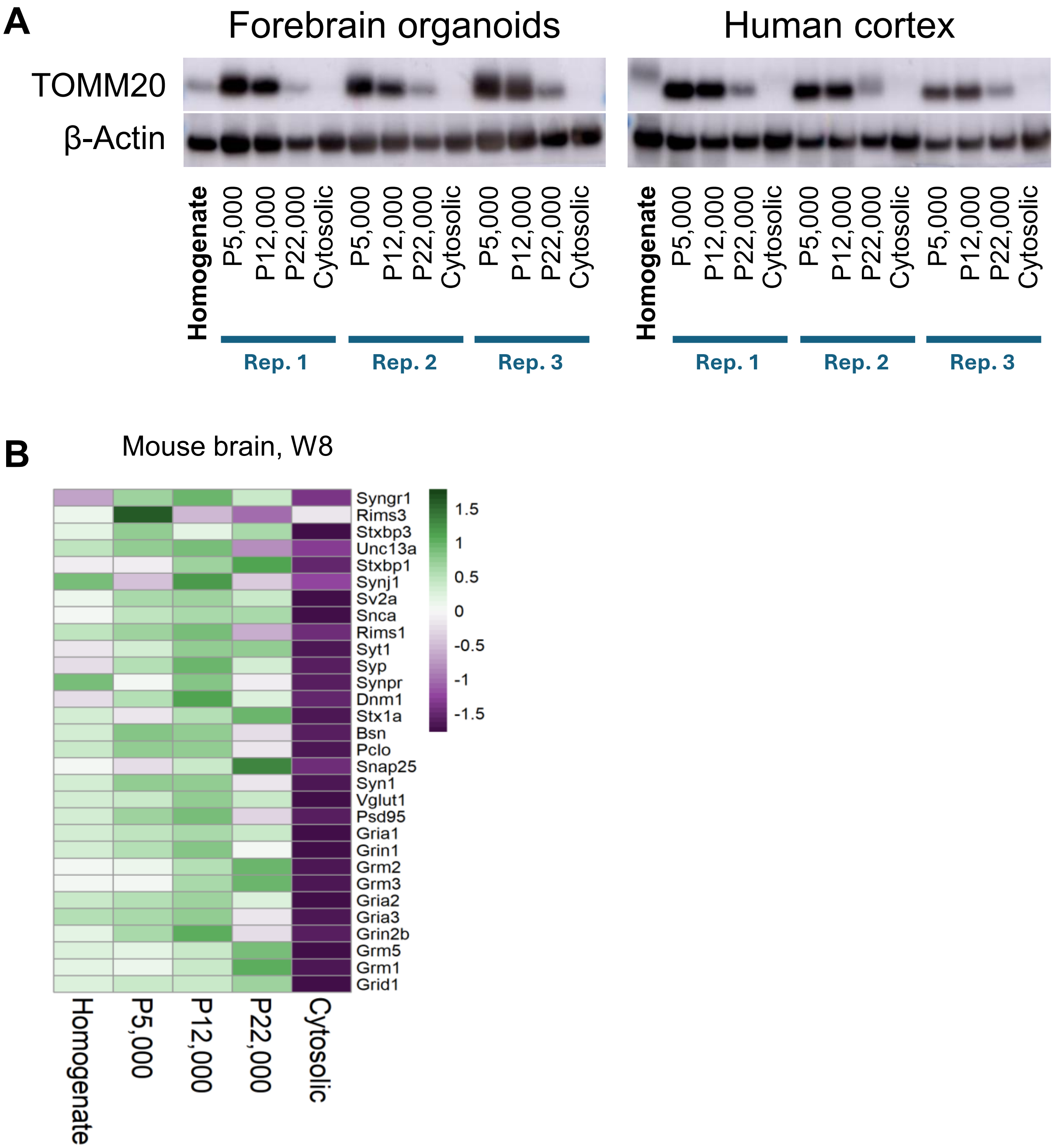


**Figure S3: A)** Western blot of the mitochondrial marker TOMM20 in differential centrifugation fractions from forebrain organoids and human cortex in three technical replicates of the differential centrifugation. **B)** Heatmap of the normalized abundances of 30 selected synaptic markers for each fraction from the differential centrifugation workflow applied to adult mouse (W8) brain tissue.

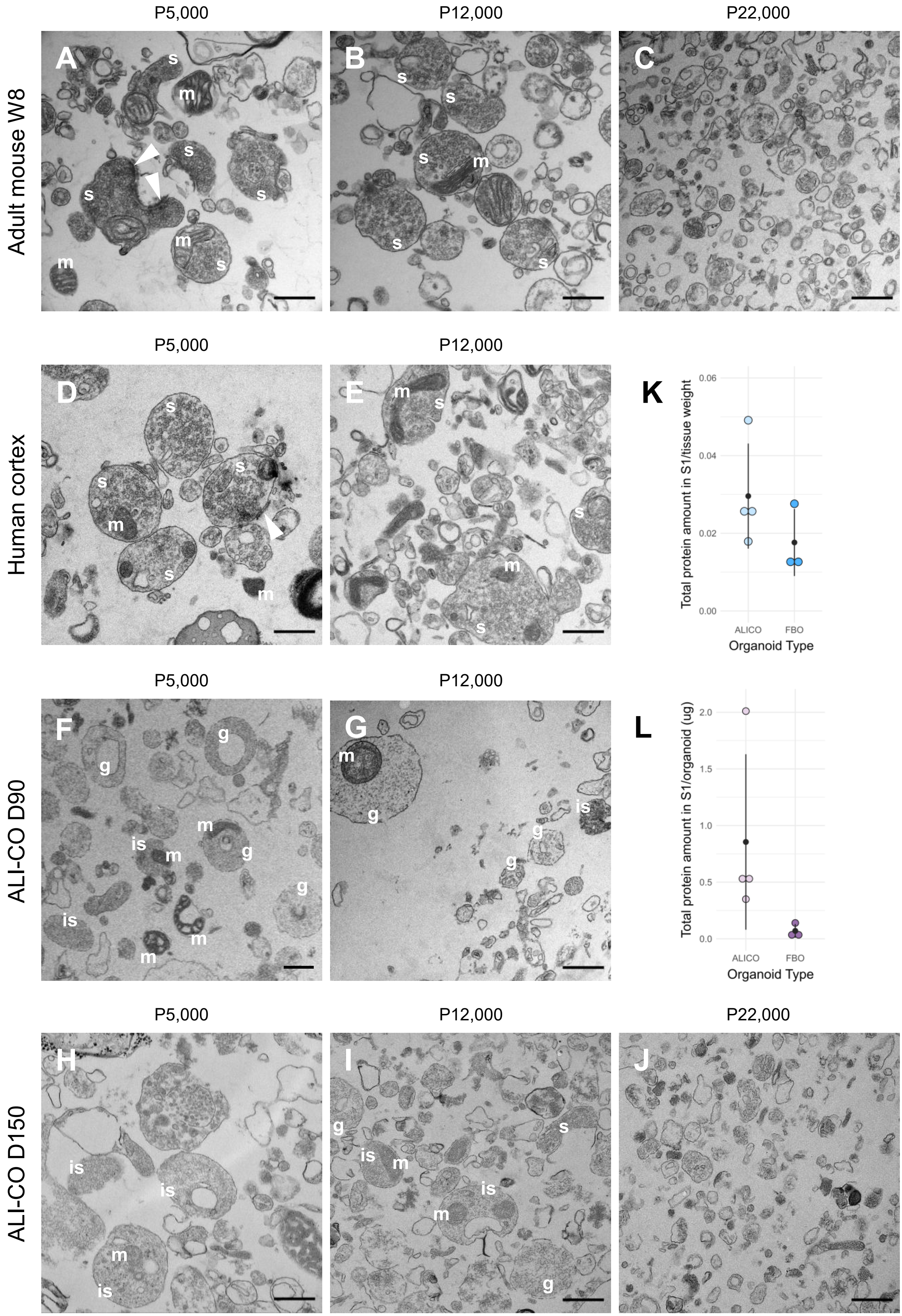

P5,000
P12,000
P22,000
Adult mouse W8
Human cortex
ALI-CO D90
ALI-CO D150
A
B
C
D
E
F
G
H
I
J
K
L
s
m
is
g
Total protein amount in S1/tissue weight
0.00
0.02
0.04
0.06
ALICO
FBO
Organoid Type
Total protein amount in S1/organoid (ug)
0.0
0.5
1.0
1.5
2.0

**Figure S4: A-J)** TEM images of differential centrifugation fractions from adult mouse, adult human cortex and ALI-COs day 90 and 150. **A-C)** Adult mouse fractions P5,000, P12,000 and P22,000. Synaptosomes (s) are clearly present in the P5,000 and P12,000 fractions. The P22,000 fraction contains mainly smaller undefined membrane structures. **D-E)** Human cortex fractions P5,000, P12,000. Synaptosomes are clearly visible in both fractions. **F-G)** Day 90 ALI-CO fractions P5,000, P12,000. A few synaptosomes or immature synaptosomes (is) could be seen, however, many structures resembled growth cone particles (g). **H-J)** Day 150 ALI-CO fractions P5,000, P12,000 and P22,000. Immature synaptosomes are seen in the P5,000 and P12,000 fractions with the presence of mitochondria and some synaptic vesicles. Scalebars **A-J)**: 500nm. s: synaptosome, is: immature synaptosome, m: mitochondria, g: growth cone particle (GCP), white arrows: postsynaptic density attached to the synaptosome. **K-L)** Comparison of yield from the differential centrifugation between ALI-CO (slice-culture) and FBO (spherical culture) organoids. **K)** Total protein amount (µg) in the first supernatant (S1) used for differential centrifugation per total weight of the used organoid tissue and **L)** per number of organoids used to begin with. The number of ALI-COs is corrected assuming that each initial cerebral organoid generates at least 4 ALI-CO slices.

***Table S1:*** *Selected synaptic proteins that are mainly expressed in the synapse.*

| Selected **synaptic markers** | | | |
|---|---|---|---|
| Gene Symbol | Protein Accession | Gene Symbol | Protein Accession |
| BSN | Q9UPA5 | RIMS3 | Q9UJD0 |
| DNM1 | Q05193 | SNAP25 | P60880 |
| GRIA1 | P42261 | SNCA | P37840 |
| GRIA2 | P42262 | STX1A | Q16623 |
| GRIA3 | P42263 | STX1B | P61266 |
| GRID1 | Q9ULK0 | STXBP3 | O00186 |
| GRIN1 | Q05586 | SV2A | Q7L0J3 |
| GRIN2B | Q13224 | SYN1 | P17600 |
| GRM1 | Q13255 | SYNGR1 | O43759 |
| GRM2 | Q14416 | SYNJ1 | O43426 |
| GRM3 | Q14832 | SYNPR | Q8TBG9 |
| GRM5 | P41594 | SYP | P08247 |
| PCLO | Q9Y6V0 | SYT1 | P21579 |
| PSD95 | P78352 | UNC13A | Q9UPW8 |
| RIMS1 | Q86UR5 | VGLUT1 | Q9P2U7 |

***Table S2:*** *Results from Wilcoxon Rank Sum tests of synaptic marker protein levels for each fraction compared to homogenate samples (n=25-30). FC: fold change. FBO: forebrain organoid; ALI-CO: air-liquid-interface cerebral organoid; D: day; P2: postnatal day 2; W8: 8-week-old.*

| Wilcoxon Rank Sum tests of **synaptic marker** levels vs. homogenate | | | | | | | | | |
|---|---|---|---|---|---|---|---|---|---|
| Fraction | Tissue type | | | | | | | | |
| | FBO (D100) | | | ALI-CO (D90) | | | ALI-CO (D150) | | |
| | FC | P-val. | V-stat. | FC | P-val. | V-stat. | FC | P-val. | V-stat. |
| P5,000 | 1.48 | 1.86E-09 | 464 | 1.24 | 5.53E-06 | 331 | 1.39 | 2.78E-04 | 284 |
| P12,000 | 1.55 | 9.31E-10 | 465 | 1.38 | 4.58E-06 | 332 | 1.34 | 1.25E-03 | 271 |
| P22,000 | 1.58 | 5.96E-10 | 442 | 1.50 | 1.24E-04 | 311 | 1.49 | 2.98E-07 | 320 |
| Cytosolic | 0.46 | 1.86E-10 | 1 | 0.36 | 2.98E-08 | 1 | 0.43 | 2.98E-08 | 0 |
| Fraction | Tissue type | | | | | | | | |
| | Human cortex | | | Mouse (P2) | | | Mouse (W8) | | |
| | FC | P-val. | V-stat. | FC | P-val. | V-stat. | FC | P-val. | V-stat. |
| P5,000 | 1.43 | 2.21E-06 | 434 | 1.26 | 1.38E-06 | 437 | 1.14 | 7.91E-04 | 381 |
| P12,000 | 1.39 | 6.52E-09 | 461 | 1.51 | 5.12E-08 | 454 | 1.32 | 1.77E-08 | 458 |
| P22,000 | 1.17 | 0.04 | 318 | 1.67 | 6.09E-06 | 427 | 1.08 | 0.24 | 268 |
| Cytosolic | 0.43 | 9.31E-10 | 0 | 0.50 | 9.31E-09 | 5 | 0.33 | 9.31E-10 | 0 |
| Fraction | Tissue type | | | | | | | | |
| | Mouse Percoll | | | | | | | | |
| | FC | P-val. | V-stat. | | | | | | |
| F3 | 1.52 | 1.30E-08 | 459 | | | | | | |
| F4 | 1.43 | 7.08E-05 | 418 | | | | | | |

*Table S3: Selected synaptic proteins that are specific for growth cones or growth cone particles (GCPs).*

| Selected **growth cone markers** | | | |
|---|---|---|---|
| Gene Symbol | Protein Accession | Gene Symbol | Protein Accession |
| AP2A1 | O95782 | MAPT | P10636 |
| ARHGDIA | P52565 | MARCKSL1 | P49006 |
| ATP1A1 | P05023 | NRXN1 | Q9ULB1 |
| CADM1 | Q9BY67 | PACS1 | Q6VY07 |
| CAMKV | Q8NCB2 | PAK1 | Q13153 |
| CANX | P27824 | PICALM | Q13492 |
| CTNNA1 | P35221 | PPFIA1 | Q13136 |
| DPYSL2 | Q16555 | PPFIA2 | O75334 |
| DSTN | P60981 | PPP2CA | P67775 |
| EXOC8 | Q8IYI6 | PTPRD | P23468 |
| GAP43 | P17677 | PTPRS | Q13332 |
| GNB2 | P62879 | SH3PXD2A | Q5TCZ1 |
| MACF1 | O94854 | SNAP29 | O95721 |
| MAP1B | P46821 | STMN1 | P16949 |

*Table S4: Results from Wilcoxon Rank Sum tests of growth cone marker protein levels for each fraction compared to homogenate samples (n=27-29). FC: fold change. FBO: forebrain organoid; ALI-CO: air-liquid-interface cerebral organoid; D: day; P2: postnatal day 2.*

| Wilcoxon Rank Sum tests of **growth cone protein** levels vs. homogenate | | | | | | | | | |
|---|---|---|---|---|---|---|---|---|---|
| Fraction | Tissue type | | | | | | | | |
| | FBO (D100) | | | ALI-CO (D90) | | | ALI-CO (D150) | | |
| | FC | P-val. | V-stat. | FC | P-val. | V-stat. | FC | P-val. | V-stat. |
| P5,000 | 1.285 | 4.70E-06 | 379 | 1.229 | 9.23E-04 | 362 | 1.35 | 3.16E-05 | 366 |
| P12,000 | 1.413 | 2.84E-06 | 382 | 1.457 | 7.10E-06 | 470 | 1.47 | 1.38E-06 | 386 |
| P22,000 | 1.304 | 0.00327 | 320 | 1.458 | 7.97E-04 | 364 | 1.46 | 2.11E-05 | 369 |
| Cytosolic | 0.536 | 2.05E-07 | 11 | 0.500 | 1.99E-05 | 27 | 0.75 | 0.00551 | 93 |

| Fraction | Tissue type | | |
|---|---|---|---|
| | Mouse (P2) | | |
| | FC | P-val. | V-stat. |
| P5,000 | 1.129 | 4.80E-05 | 340 |
| P12,000 | 1.259 | 2.76E-06 | 358 |
| P22,000 | 1.424 | 0.00273 | 302 |
| Cytosolic | 0.627 | 1.26E-06 | 16 |
